# A Practical Preprocessing Pipeline for Concurrent TMS-iEEG: Critical Steps and Methodological Considerations

**DOI:** 10.1101/2025.08.13.670238

**Authors:** Zhuoran Li, Xianqing Liu, Joshua Tatz, Umair Hassan, Jeffrey B Wang, Corey J. Keller, Nicholas T. Trapp, Aaron D. Boes, Jing Jiang

## Abstract

Transcranial magnetic stimulation combined with intracranial EEG (TMS-iEEG) has emerged as a powerful approach for probing the causal organization and dynamics of the human brain. Despite its promise, the presence of TMS-induced artifacts poses significant challenges for accurately characterizing and interpreting evoked neural responses. In this study, we present a practical preprocessing pipeline for single pulse TMS-iEEG data, incorporating key steps of re-referencing, filtering, artifact interpolation, and detrending. Using both real and simulated data, we systematically evaluated the effects of each step and compared alternative methodological choices. Our results demonstrate that this pipeline effectively attenuated various types of artifacts and noise, yielding cleaner signals for the subsequent analysis of intracranial TMS-evoked potentials (iTEPs). Moreover, we showed that methodological choices can substantially influence iTEPs outcomes. In particular, referencing methods might strongly affect iTEP morphology and amplitude, underscoring the importance of tailoring the referencing strategy to specific signal characteristics and research objectives. For filtering, we recommend a segment-based strategy, i.e., applying filters to data segments excluding the artifact window, to minimize distortion from abrupt TMS-related transients. Overall, this work represents an important step toward establishing a general preprocessing framework for TMS-iEEG data. We hope it encourages broader adoption and methodological development in concurrent TMS-iEEG research, ultimately advancing our understanding of brain organization and TMS mechanisms.

**Highlights:**

1. We presented a practical preprocessing pipeline for single pulse TMS-iEEG data, incorporating key steps of re-referencing, filtering, artifact interpolation, and detrending.
2. The pipeline effectively attenuates multiple types of artifacts and noise, enabling accurate characterization of evoked neural responses.
3. Methodological alternatives for each preprocessing step were evaluated using real and/or simulated datasets.
4. Re-referencing substantially affects the morphology and amplitude of intracranial TMS-evoked potentials and requires careful consideration.
5. A segment-based filtering strategy is recommended to better minimize distortion from TMS-related artifacts.

## Introduction

Recent advances in combining brain stimulation techniques, e.g., transcranial magnetic stimulation (TMS), with concurrent neural measurement techniques, e.g., functional MRI (fMRI) or scalp electroencephalography (EEG), have opened new avenues to probe the causal organization and dynamics of the human brain. Among these, concurrent TMS with intracranial EEG (iEEG) has emerged as a novel and powerful approach, enabling direct measurement of the immediate neural consequences of stimulation with both high temporal and spatial resolutions. Prior studies demonstrated that single-pulse TMS (spTMS) applied to cortical regions can elicit intracranial TMS-evoked potentials (iTEPs) not only near the stimulation site (Wang et al., 2024), but also in anatomically distant yet functionally connected areas (Li et al., 2025; Solomon et al., 2025; Solomon et al., 2024; Trapp et al., 2025; Wang et al., 2024). For example, TMS applied to the dorsolateral prefrontal cortex (DLPFC) elicited significant iTEPs in the subgenual anterior cingulate cortex (sgACC) (Trapp et al., 2025), supporting the hypothesis that sgACC is a key downstream target underlying the antidepressant effects of DLPFC TMS (Oathes et al., 2023; Siddiqi and Fox, 2024). These findings highlight TMS-iEEG as a valuable tool for mapping causal brain networks and elucidating the mechanisms by which cortical stimulation modulates activity and functions in distant regions.

Despite its promise, TMS-iEEG data are often contaminated by multiple sources of artifacts (Figure 1A–B), including stimulation-related artifacts, e.g., amplifier saturation, TMS-induced ringing, and decay artifacts, as well as conventional iEEG artifacts such as line noise, motion artifacts, and, in patients with epilepsy, epileptiform discharges (Mercier et al., 2022). Critically, many of these stimulation and iEEG artifacts overlap with the early post-TMS period, typically tens to hundreds of milliseconds after stimulation, when meaningful TMS-evoked neural responses are expected to emerge (Hassan et al., 2025; Li et al., 2025; Solomon et al., 2024; Wang et al., 2024). As such, an effective and well-designed preprocessing pipeline is essential for attenuating these confounds and accurately characterizing the temporal dynamics of iTEPs.

**Figure 1.**
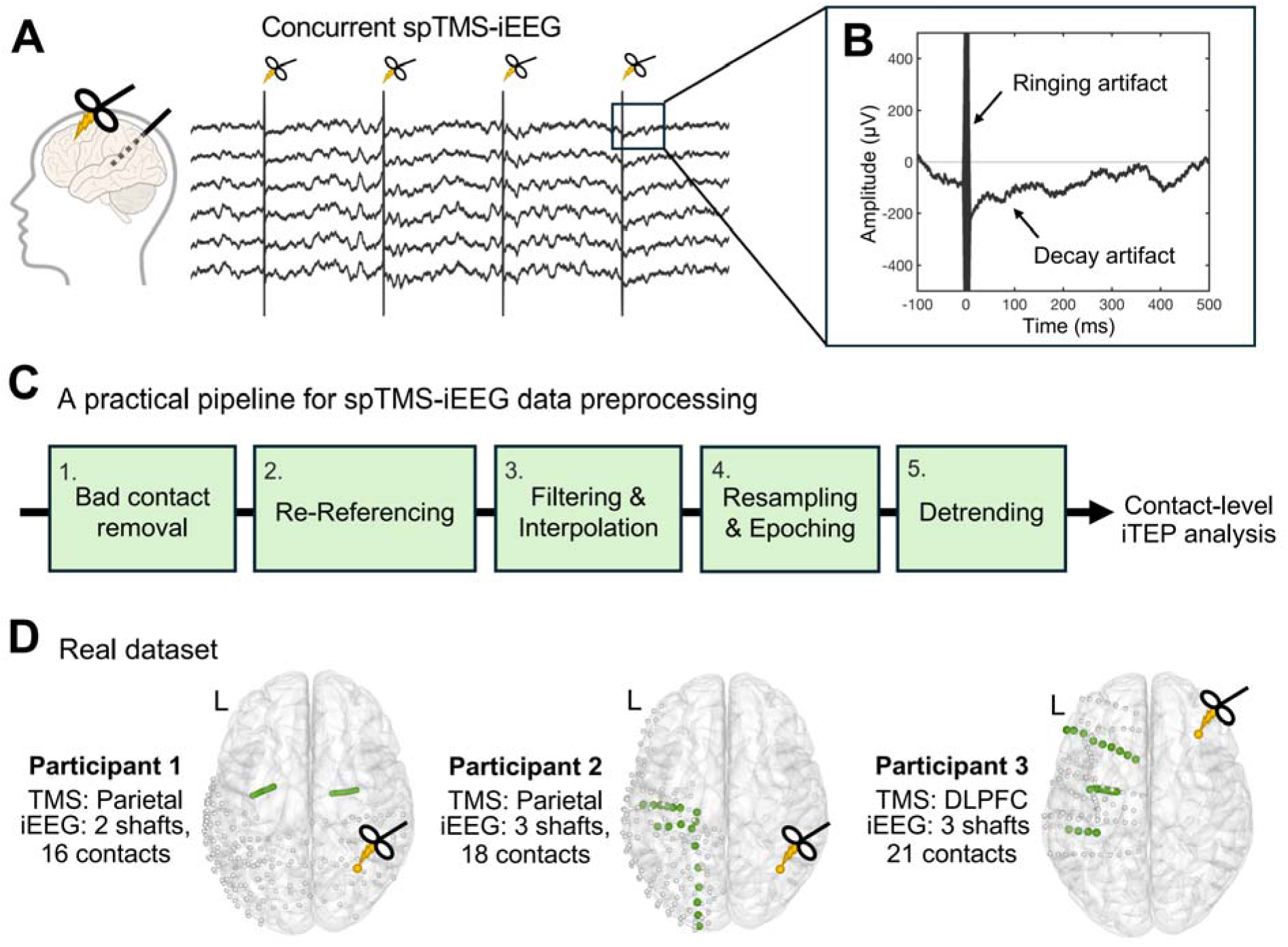
Overview of spTMS-iEEG Protocol, Artifacts, Preprocessing Pipeline, and Dataset. (A) Schematic of the concurrent spTMS-iEEG protocol. (B) A representative example trial illustrating artifacts and noise observed in raw spTMS-iEEG data. TMS-induced large-amplitude ringing artifacts, and long-lasting decay artifacts were marked. (C) A practical preprocessing pipeline for spTMS-iEEG data, incorporating critical steps of re-referencing, filtering, artifact interpolation, and detrending. (D) Overview of the dataset used to evaluate the preprocessing pipeline. Concurrent spTMS-iEEG data were collected from three participants. Depth electrode shafts (55 contacts in total) targeting key regions of interest (e.g., hippocampus, amygdala, and subgenual anterior cingulate cortex; see Methods and Supplementary Methods) are highlighted and used to systematically assess the preprocessing pipeline.

Existing TMS-iEEG studies (Li et al., 2025; Solomon et al., 2025; Solomon et al., 2024; Trapp et al., 2025; Trapp et al., 2024; Wang et al., 2024) have adopted preprocessing pipelines that typically include several key steps: re-referencing, artifact interpolation, bandpass and notch filtering, and detrending. These steps are designed to remove various artifacts or noise in TMS-iEEG data. Specifically, re-referencing aims to remove noise shared across electrode contacts; interpolation aims to correct high-amplitude artifacts surrounding the TMS pulse; filtering aims to attenuate undesired spectral components such as line noise, low-frequency drift, and high-frequency noise; and detrending algorithms aim to reduce post-stimulation decay artifacts. However, these studies differ considerably in both the inclusion and implementation of these steps. Moreover, the impact of each preprocessing step and the difference between methodological alternatives remain underexplored. Given that TMS-iEEG is still in its early stages, with the first study in humans published by our group in 2024 (Wang et al., 2024), a systematic evaluation of these preprocessing steps is warranted. Such efforts will provide guidance for future TMS-iEEG studies and contribute to the development of standardized, replicable pipelines.

In this study, we present a practical preprocessing pipeline for spTMS-iEEG data incorporating these key steps. Using real and simulated data, we systematically quantify the impact of each step and compare alternative methodological choices and parameters. Together, this work offers an important first step toward informing optimal preprocessing frameworks for TMS-iEEG. In turn, we hope this promotes broader adoption and methodological development of TMS-iEEG among researchers and clinicians.

## Materials and Methods

### Real Dataset

We analyzed data from three neurosurgical patients (2 female, ages 23 and 24; 1 male, age 34). These patients were with medically intractable epilepsy and admitted to the University of Iowa Hospitals and Clinics with intracranial electrode implantation to localize their seizure focus. All experimental procedures were approved by the University of Iowa Institutional Review Board (IRB), with written informed consent obtained from all patients.

During the concurrent spTMS-iEEG experiment, 50 single pulses of TMS were delivered at a frequency of 0.5 Hz over either DLPFC or parietal cortex, with stimulation intensity set at 120% of resting motor threshold (MT), or 100% MT if 120% was not tolerated. iEEG data were simultaneously recorded using a multichannel data acquisition system (ATLAS, Neuralynx, Tucson, AZ), with a reference electrode placed in patients’ subgaleal space. All three patients were implanted with depth electrodes (Stereo-electroencephalography, sEEG) and/or subdural grid/strip arrays (electrocorticography, ECoG) (Ad-Tech Medical, Racine, WI). Demographic and experimental details for each patient are provided in Supplementary Methods and Table S1.

To clarify terminology in this study, the term “electrode” refers to the physical device implanted in the brain; the term “contact” refers to the individual recording sites along the electrode and their corresponding recorded signals (alternatively referred to as “channel” in some literature) (Parish et al., 2023). This study mainly focused on sEEG contacts, given the growing clinical adoption of sEEG and its unique advantage in recording TMS-evoked responses from deep brain structures over ECoG and non-invasive modalities such as scalp EEG (Unnwongse et al., 2023). We specifically selected sEEG shafts covering the deep brain regions of interest (ROIs), including the hippocampus, amygdala, and sgACC, yielding a total of 55 sEEG contacts across three participants. Additional methodological details, including ROI atlases, contact localization and selection, etc., are further described in Supplementary Methods.

### Overview of Preprocessing Pipelines

The overview of our preprocessing pipeline is illustrated in Figure 1C. The workflow comprises several steps: (1) removal of bad contacts, (2) re-referencing, (3) filtering and artifact interpolation, (4) resampling and epoching, and (5) detrending. All preprocessing procedures were implemented using custom MATLAB scripts (R2024b) and the FieldTrip toolbox (Oostenveld et al., 2011). Detailed descriptions of each step are provided below.

## 1. Bad Contact Removal

The first preprocessing step is identifying and removing “bad” contacts based on clinical reports and experimental documentation. Specifically, contacts located within the seizure onset zone, typically identified by clinicians based on the presence of epileptiform spikes or high-frequency oscillations, and those exhibiting excessive noise due to non-neuronal signals, including faulty sensors, bad connections, etc. (Hattab et al., 2024), should be removed. This step is critical to avoid misleading conclusions from problematic contact and to prevent noise contamination during subsequent preprocessing steps, e.g., re-referencing.

## 2. Re-Referencing

Re-referencing aims to reduce contamination from the reference site, such as signals from volume conduction or physiological artifacts (e.g., muscle activity) (Li et al., 2018; Michelmann et al., 2018). We evaluated monopolar referencing method and two typical re-referencing methods, i.e., bipolar, and common-average methods. The monopolar method retains the raw recording signals referenced to the designated reference electrode, which preserves all activity but is vulnerable to reference noise. The bipolar re-referencing creates a second spatial derivative signal by subtracting the data from its closest neighbor within the same shaft. It can enhance detection of focal activity and suppress shared noise, but may attenuate broader, network-level signals (Parish et al., 2023). The common-average re-referencing subtracts the mean signal across all contacts. It can suppress global artifacts while preserving localized signals but may be biased by noisy or high-amplitude contacts (Huang et al., 2024). We applied all three methods to real spTMS-iEEG data to examine their effects on the waveform morphology and amplitude of the resulting iTEPs.

Furthermore, given the difficulty to distinguish the true signal and noise in real recordings, we conducted simulations to quantitatively evaluate the performance of these three referencing methods (Figure 2B). We simulated a four-contact sEEG shaft model, following procedures adapted from a previously published study (Michelmann et al., 2018). First, three latent sources were defined using three 1-sec segments of real spTMS-iEEG data and designated as source 1 (S1), source 2 (S2), and reference source (R). All signals were demeaned and scaled to unit variance, with R further scaled to 10% of the amplitude of S1 and S2. Then, we constructed a linear mixture model simulating recordings from four contacts in a shaft (C1–C4), where S1 and S2 were placed on C1 and C4, respectively, and spread to neighboring contacts with strengths of 1/a, 1/a^2^, and 1/a^3^. The parameter a was varied from 10 to 1.02. Each signal was then referenced to R with a fixed coefficient of –1. Gaussian pink noise was also added to each contact, with variance varying from 0% to 100% of the source variance. Next, the three referencing methods were applied to the simulated data. Last, performance was evaluated using sensitivity (mean Pearson correlation with the close, corresponding source), specificity (1 minus the correlation with the distant source), and a combined metric (sensitivity × specificity). Each condition was repeated 1,000 times to ensure robust estimates. Further details of the simulation procedure, including the selected real data segments, parameter specifications, etc., are described in the Supplementary Methods.

**Figure 2.**
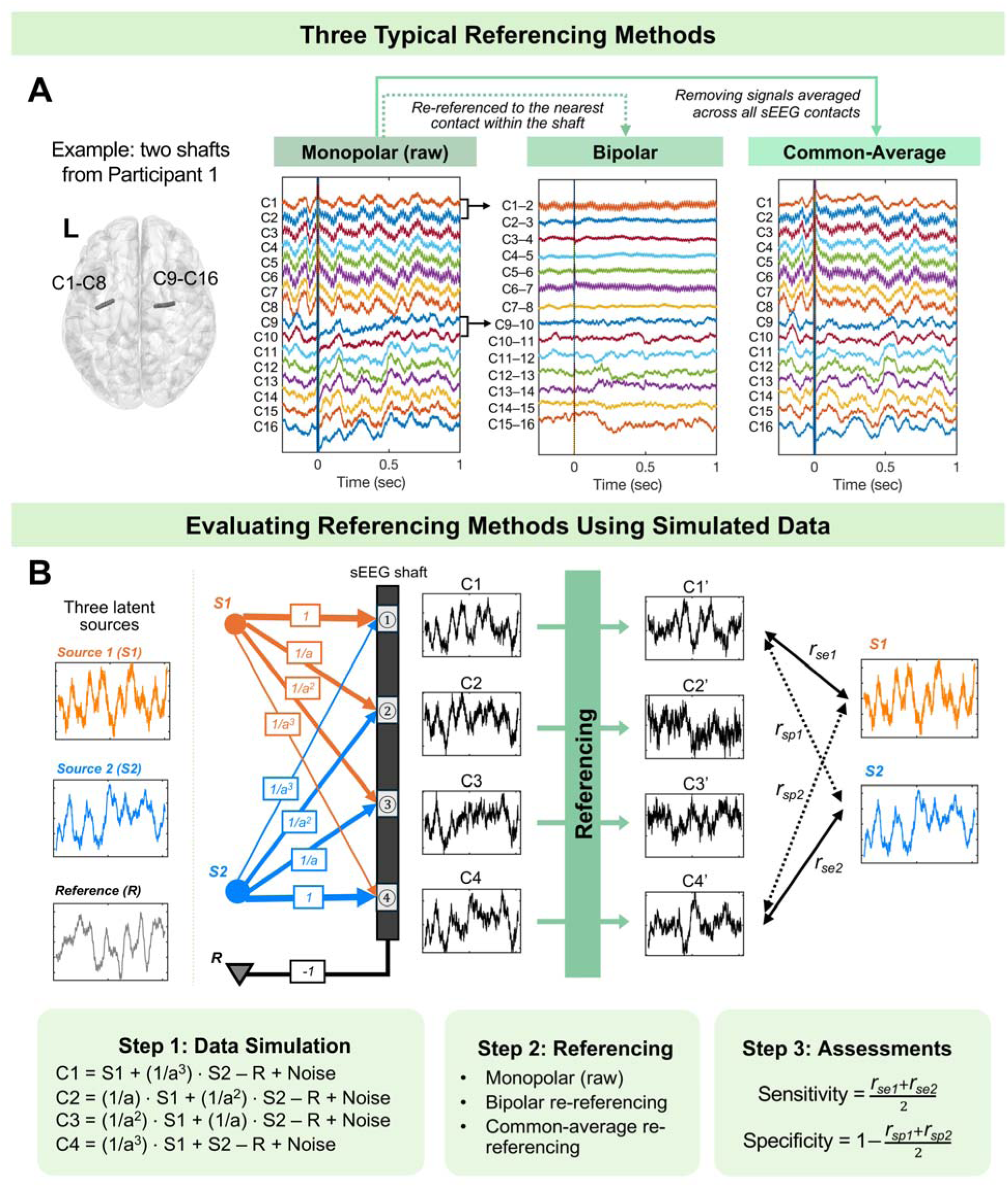
Typical Referencing Methods and Workflow of Data Simulation. (A) Schematic illustration of three typical referencing methods: monopolar, bipolar, and common-average methods. (B) Workflow of the data simulation for evaluating the performance of three referencing methods. A linear mixture model was constructed to simulate iEEG signals of a four-contact sEEG shaft, using two hidden sources (S1, S2) and one reference source (R), each from real iEEG data. S1 and S2 were assigned to contacts at the two ends of the shaft and allowed to influence adjacent contacts with decreasing weights of 1/a, 1/a^2^, and 1/a^3^, where a reflects the spatial spread of the signal. The mixed data were further added with pink noise and then subjected to one of the three referencing methods. The performance of referencing methods was quantified by analyzing the sensitivity and specificity: Sensitivity was computed as the mean Pearson correlation between the re-referenced signal at the two ends of the shaft and its nearby true source; Specificity was computed as 1 minus the mean Pearson correlation with the opposite, distant source.

## 3. Filtering and Interpolation

This preprocessing step consists of two parts: frequency filtering and interpolation of TMS-related artifacts. For frequency filtering, we applied a notch filter (57–63 Hz and seven harmonics, third-order Butterworth; Fieldtrip toolbox) and a zero-phase bandpass filter (third-order Butterworth bandpass filter; Fieldtrip toolbox). Two bandpass filtering settings were tested: 1–200 Hz and 2–200 Hz. The high cutoff of 200 Hz was chosen to preserve broadband neural oscillations while attenuating higher-frequency fluctuations, which are typically considered pathological or non-physiological (Buzsáki et al., 2012). The low cutoff (1 or 2 Hz) was selected to exceed the stimulation frequency (0.5 Hz), minimizing contamination from stimulation-related periodicity while remaining low enough to preserve meaningful low-frequency neural signals and avoid distortion around strong evoked responses (Zhang et al., 2024). As 1-and 2-Hz low cutoffs have been used in different prior TMS-iEEG studies (Li et al., 2025; Solomon et al., 2024; Trapp et al., 2025; Wang et al., 2024), we included and compared both.

Interpolation aims to recover the signal within the time window contaminated by strong artifacts surrounding the TMS pulse. These artifacts typically last only a few milliseconds but can be prolonged due to electromagnetic decay within the electrode system or filtering effects of the amplifier (Hernandez-Pavon et al., 2023). Based on inspection of the raw recordings (Figure S1), we defined the artifact window as [–10, +15 ms]. To preserve adjacent signal integrity, we interpolated this window using surrounding iEEG data. Specifically, equal-length segments immediately before and after the artifact window are reversed, tapered (e.g., linearly from 1 to 0 and 0 to 1, respectively), and summed to synthesize replacement data with a similar spectral profile (Li et al., 2025; Solomon et al., 2024; Trapp et al., 2025; Wang et al., 2024).

To prevent the impact of large TMS-related transients on filtering (Hernandez-Pavon et al., 2022; Rogasch et al., 2017), a typical strategy is to interpolate the artifact window before applying filters to the entire data (referred to as “full-length filtering strategy”). However, this strategy might be suboptimal for spTMS-iEEG data, as interpolation does not fully resolve the sharp transition between the pre-TMS signal (typically near zero) and the post-TMS signal (which may contain large evoked responses or decay-related artifacts). These discontinuities can introduce spurious fluctuations during filtering. To address this, we employed a segment-based filtering strategy, in which the artifact window was first removed, and the filters (both notch and bandpass filters) were applied only to the remaining segments individually (Figure 4A). Interpolation was performed afterward. By applying filters only to relatively clean segments without strong artifacts or data transition, this strategy might reduce the risk of distortion and yield more reliable filtering performance.

In this study, we tested both two filtering strategies (segment-based vs. full-length-based) and with two bandpass settings (1–200 Hz vs. 2–200 Hz). To quantitatively assess and compare the filtering-induced distortions under these conditions, we analyzed the signal magnitude at the time points immediately beyond the onset (–11 ms) and offset (+16 ms) of this TMS artifact window. This analysis was conducted at both the trial and contact levels. At the trial level, we calculated the proportion of trials whose magnitude increased while retaining the same polarity after filtering. A lower proportion indicates fewer filtering-induced distortions. At the contact level, we calculated the average magnitude across trials for each contact and performed paired-sample t-tests to compare signal magnitudes before and after applying each filtering method.

## 4. Resampling and Epoching

After filtering and interpolation, the data were resampled to 1000 Hz to reduce file size and improve computational speed in subsequent analysis. Resampling is recommended after artifact interpolation to prevent smearing of strong TMS artifacts into neighboring time points. The continuous data were then epoched into segments from –250 to +500 ms relative to each TMS pulse onset to facilitate subsequent trial-based computations and further analysis of iTEPs.

## 5. Detrending

We implemented an adaptive detrending algorithm (ADA) (Casula et al., 2017) to detect and eliminate potential decay artifacts. The epoched iEEG signals within the 16– 500 ms interval were modeled for each contact and trial, using either a linear regression or a two-exponential function. The parameters [a, b] for the linear model were estimated using weighted linear least squares, while the parameters [A_1_, a_1_, A_2_, a_2_] for the exponential model were estimated using weighted nonlinear least squares. In both cases, weights were set optimally, proportional to the inverse of the trial-wise variance.

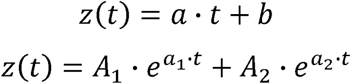

If neither model returned a robust fit, it indicated the absence of a prominent linear or exponential decay trend. If fitting was successful, the model with the lower Akaike Information Criterion (AIC) was selected, and the corresponding fitted trend was then subtracted from the signal. It is worth noting that fitting either model could produce edge discontinuity at the end of the epoched time interval (i.e., 500 ms). This was not a concern in our pipeline, as detrending was performed as the final preprocessing step and iTEP analyses were restricted to 16–500 ms window. However, if subsequent computations require smooth signal transitions beyond this interval, such as spectral analysis, or if detrending is performed before filtering in alternative pipelines, a cosine taper can be applied to the fitted trend to minimize its influence at the later timepoints. In such cases, an additional interpolation step is further needed to update the data within the artifact window.

To quantitatively assess the effects of ADA in removing the decay trends, we analyzed signal magnitude within the 16–500 ms post-TMS interval both before and after applying ADA, at the trial and contact levels. At the trial level, we calculated and plotted time series of the signal magnitude before and after applying ADA, for both segment-based and full-length-based filtered data. This enabled us to visually identify the time window where ADA most effectively reduced decay artifacts. At the contact level, we first computed the area under the curve (AUC) within this decay window, and averaged across trials for each contact. Paired-sample t-tests were then conducted to compare the AUC across different conditions. A lower level of AUC reflected a cleaner signal with less decay artifacts.

## 6. Contact-level iTEP Analysis

We next evaluated the overall effects of preprocessing on the contact-level iTEPs, defined as the average response across trials for each contact. To ensure a reliable iTEP estimation, trials exhibiting abnormally large amplitudes, indicative of residual TMS-related artifacts, interictal epileptiform activity, or other non-physiological noise, were identified and excluded. Following strategies from iEEG studies combined with intracranial electrical stimulation (iES) (Sawada et al., 2022; van Blooijs et al., 2023) and prior TMS-iEEG studies (Li et al., 2025; Trapp et al., 2025), trials were rejected if they met any of the following criteria: (1) signal at 16 ms post-TMS exceeded 10 standard deviations (SD) from the pre-stimulation baseline (–250 to –50 ms) or an absolute threshold of 200 μV; (2) signal exceeded 50 baseline SDs or 300 μV at any time point in the epoch. These threshold parameters were chosen to balance sensitivity to artifacts and retention of physiologically valid responses, but may require adjustment based on specific dataset characteristics, e.g., target region, stimulation protocol, and iEEG acquisition system. Contacts with insufficient retained trials (e.g., < 50% of total trials) were considered too noisy and excluded from further analysis. For remaining contacts, trials were averaged and baseline-corrected by subtracting the mean amplitude of the pre-stimulation baseline period, yielding the final contact-level iTEPs.

## Results

### 1. Simulation-Based Performance of Various Referencing Methods

To quantitively compare the performance of different referencing methods, we simulated a four-contact sEEG shaft and evaluated sensitivity (similarity to the true source) and specificity (dissimilarity from a distant source). The monopolar method exhibited high sensitivity and specificity across a broad range of conditions, remaining robust until noise exceeded 0.5 and 1/a^3^ exceeded ∼0.8 (Figure 3, Figure S2). In contrast, bipolar and common-average methods exhibited high sensitivity only when the noise ratio and 1/a^3^ were both low, with a marked decline as either parameter increased. Specificity was highest with the bipolar method even under high noise or large 1/a^3^ conditions, while the common-average method overall showed the lowest specificity.

**Figure 3.**
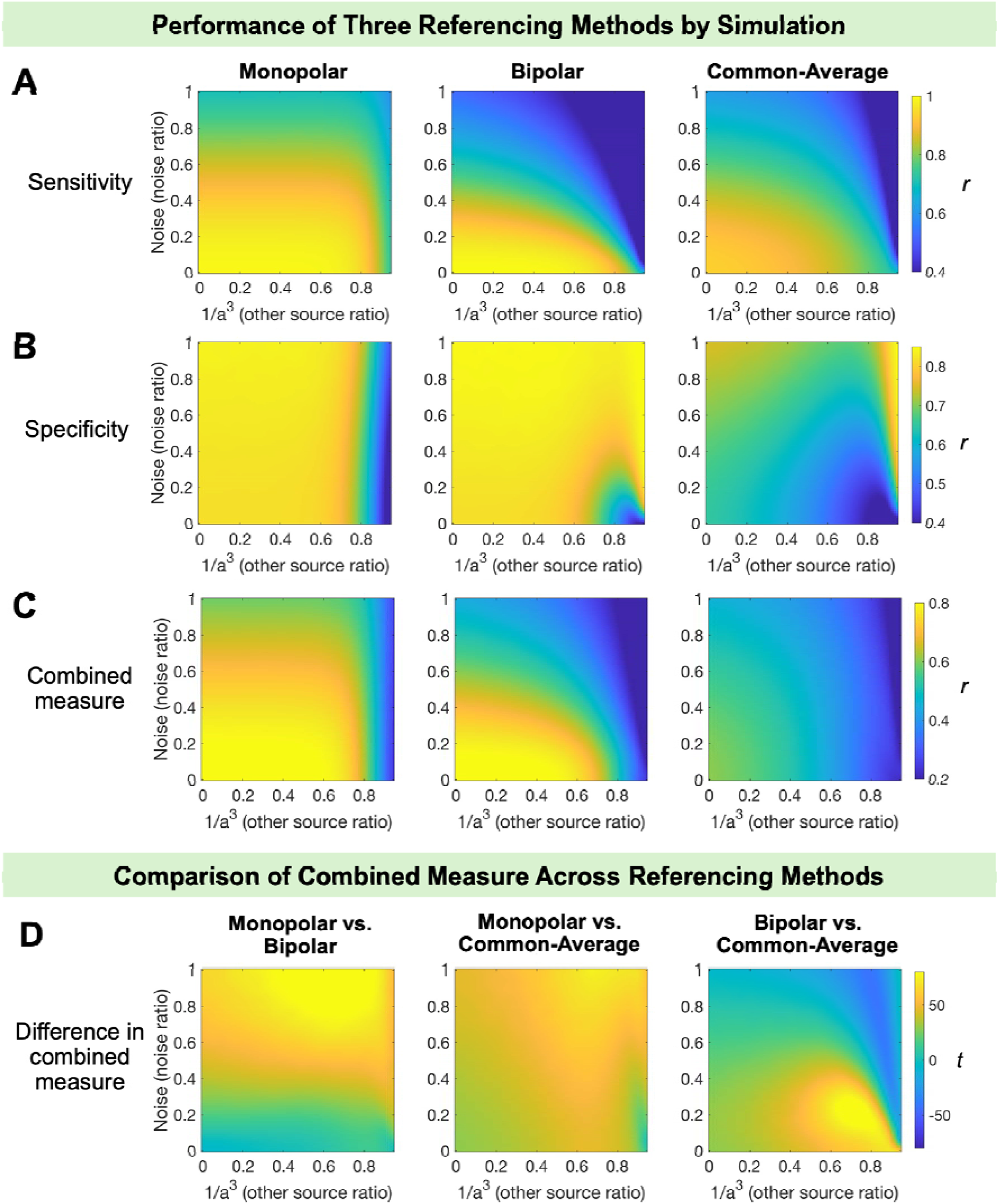
Simulation-Based Performance of Three Referencing Methods. Simulated data (N = 1,000) were used to evaluate the performance of three referencing methods in terms of sensitivity and specificity. Sensitivity was quantified as the Pearson correlation between the referenced signal and the true source, while specificity was defined as 1 minus the correlation with an interfering source. A combined measure was computed by multiplying sensitivity and specificity (see Methods for more information about the simulation and computation). (A) Sensitivity of three referencing methods. The monopolar method maintained a high sensitivity across a wide range of conditions, while sensitivity for bipolar and common-average methods declined substantially with a higher noise or 1/a^3^. (B) Specificity of three referencing methods. Both monopolar and bipolar methods exhibited high and relatively stable specificity across most conditions. The common-average method exhibited an overall low specificity. (C) Combined measure of three referencing methods. The monopolar method maintained a high performance across most conditions. Bipolar reference performed well only when both noise and 1/a^3^ were low. The common-average method exhibited an overall low performance. (D) Difference of the combined measure across three methods. The t-values were estimated based on the 1,000 times of stimulation. The monopolar method overall outperformed both the bipolar and common-average methods.

Using a combined measure (sensitivity × specificity), the monopolar method exhibited high performance across most conditions (Figure 3C), overall surpassing the bipolar and common-average method (Figure 3D). The bipolar method performed well with a low-to-moderate 1/a^3^ but declined with increasing noise, whereas the common-average method exhibited a relatively low performance across most conditions. It should be acknowledged that this suboptimal performance of common-average method may stem from the limited number of simulated contacts, which may be insufficient for accurately estimating common noise (see Discussion for further context) and thus may not fully reflect performance in real datasets that typically include dozens to hundreds of contacts.

To assess how much these results were driven by the relatively weak reference source chosen (10% of the target source amplitude), we conducted additional simulations varying the reference amplitude to 20%, 50%, and 100% of the target sources. Results showed that as reference signal became stronger, the performance of monopolar method declined and ultimately fell below that of the bipolar method (Figure S3). The bipolar and common-average performance remained relatively stable. In particular, the bipolar method maintained high performance when both noise and 1/a^3^ were low, even at the strongest reference amplitude (100% variance level).

Together, our simulations suggest that the optimal referencing methods might depend on specific signal characteristics. When the designated reference site is minimally contaminated, as reflected by low signal similarity across contacts in real recordings, the monopolar method provides a robust and superior performance. In contrast, when a strong shared, non-target activity is observed, re-referencing becomes increasingly important to better detect the true evoked response.

### 2. Effects of Different Filtering Strategies and Bandpass Settings

We evaluated the effects of filters using two filtering strategies (segment-based vs. full-length-based) and two bandpass settings (1–200 Hz vs. 2–200 Hz). Example trials (Figure 4B) showed that all filters effectively removed line noise and low-frequency drifts. However, notable differences between the two filtering strategies were observed in the –100 and +100 ms window surrounding the TMS pulse. Specifically, the segment-based strategy yielded signals closer to zero, whereas the full-length-based strategy introduced additional distortions, as evident in pre-TMS signal elevation in one example (Figure 4B, right). Regarding the frequency setting of bandpass filters, 1-Hz and 2-Hz filters yielded similar results, though the 2-Hz filter slightly better suppressed low-frequency drift (Figure 4B, left).

**Figure 4.**
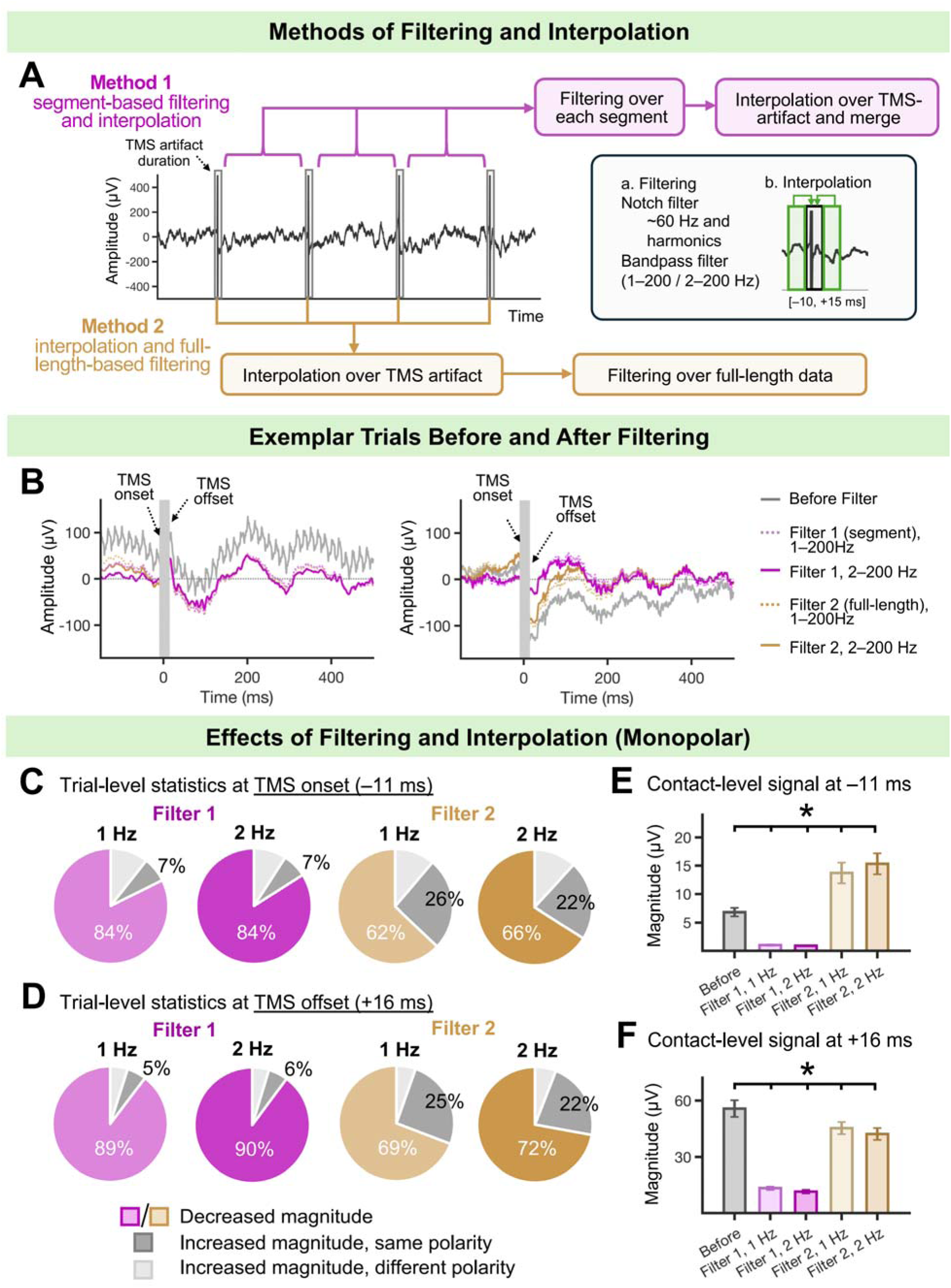
Methods and Effects of Filtering and Interpolation. (A) Schematic illustrating different methodological choices of filtering and interpolation. Two filtering strategies (i.e., segment-based and full-length-based methods) and two bandpass filters settings (i.e., 1–200 Hz and 2–200 Hz) were designed and evaluated. (B) Example trials from two contacts across two participants. All filters could remove line noise and post-TMS drift. Notably, the segment-based strategy yielded cleaner signals near the TMS onset and offset. In contrast, the full-length strategy introduced noticeable filtering-induced distortions near TMS onset, particularly in the second example (right panel). The difference between the 1-Hz and 2-Hz filters was minor. (C–F) Effects of filtering and interpolation on monopolar referenced data (see Figure S4 for results in bipolar and common-average re-referenced data). (C–D) Trial-level statistics at the TMS onset (C) and offset (D). All the trials from 55 contacts from 3 participants were analyzed. The segment-based method yielded a higher proportion of trials with decreased magnitude and a lower proportion with increased magnitude. Results for 1-Hz and 2-Hz filters were close. (E–F) Contact-level statistics at the TMS onset (E) and offset (F). Error bars represent the standard error across contacts. Here, three contacts with overwhelmingly large magnitudes at TMS offset (over three scaled mean absolute deviation from the group average) were considered as too noisy and thus excluded, resulting in a total of 52 contacts for statistical analysis. All the pairwise comparisons between conditions were significant (as marked by the bars and asterisks, p_FDR_ < .05). Overall, the segment-based filtering strategy with the 2–200 Hz filter yielded the lowest magnitude at both the TMS onset and offset.

To quantitatively assess these effects, we analyzed signal magnitudes at the onset (–11 ms) and offset (+16 ms) of the TMS artifact window at both the trial and contact levels. Trial-level analyses revealed that >80% of trials exhibited reduced magnitude following segment-based filtering, compared to ∼70% with full-length filtering, across both bandpass settings and all referencing methods (Figures 4C–D, Figure S4). Furthermore, the proportion of trials with filtering-induced distortion, defined as increased magnitude with unchanged polarity, was lower with segment-based filtering (∼10% of trials) than with full-length filtering (>20%). Differences between the 1-Hz and 2-Hz filters were minimal.

At the contact level, segment-based filtering significantly reduced signal magnitude at the TMS onset across all referencing methods and both bandpass settings (ts < –5.17, p_FDR_ < .001, Figure 4E, Figure S4). In contrast, full-length-based filtering unexpectedly increased the onset signal magnitude for monopolar and common-average referenced data (ts > 4.80, p_FDR_ < .001; Figure 4E, Figure S4), but not for bipolar referenced data (p_uncorrected_ > .37). At the TMS offset, segment-based filtering again significantly reduced signal magnitude across all conditions (ts < –3.17, p_FDR_ < .004, Figure 4F, Figure S4). Full-length filtering also led to statistically significant magnitude reductions in most conditions (ts < –2.44, p_FDR_ < .021), except for bipolar re-referenced data with 1-Hz filter, where only a similar but non-significant trend was observed (t(48) = –1.81, p_uncorrected_ = .077).

Direct comparisons further confirmed that segment-based filtering consistently yielded significantly lower magnitudes than full-length filtering at both the TMS onset and offset across all referencing methods and bandpass settings (ts < –2.71, p_FDR_ < .011). Besides, the 2-Hz filter resulted in lower magnitude than the 1-Hz filter at both the TMS onset and offset for monopolar and common-average referenced data (ts < – 2.39, p_FDR_ < .015). For the bipolar referenced data, the 2-Hz filter yielded a significantly lower magnitude at the TMS onset using the segment-based filtering (t(48) = –2.32, p_FDR_ = .026), with no significant differences observed at the TMS offset or by the full-length-based filtering (p_uncorrected_ > .10). Together, these results suggest that the segment-based filtering strategy, particularly with the implementation of a 2–200 Hz bandpass filter setting, offers a superior performance in suppressing unwarranted spectral components while minimizing filtering-induced distortions.

### 3. Effects of Detrending in Removing Long-Lasting Artifacts

We used ADA to model and remove post-TMS decay artifacts in the 16–500 ms window, across data processed with three referencing methods and two filtering strategies. The bandpass filter was fixed at 2–200 Hz, given its comparable yet slightly superior performance over 1–200 Hz. Example trials (Figure 5B) showed that ADA effectively removed the residual decay artifacts within the early post-TMS window.

**Figure 5.**
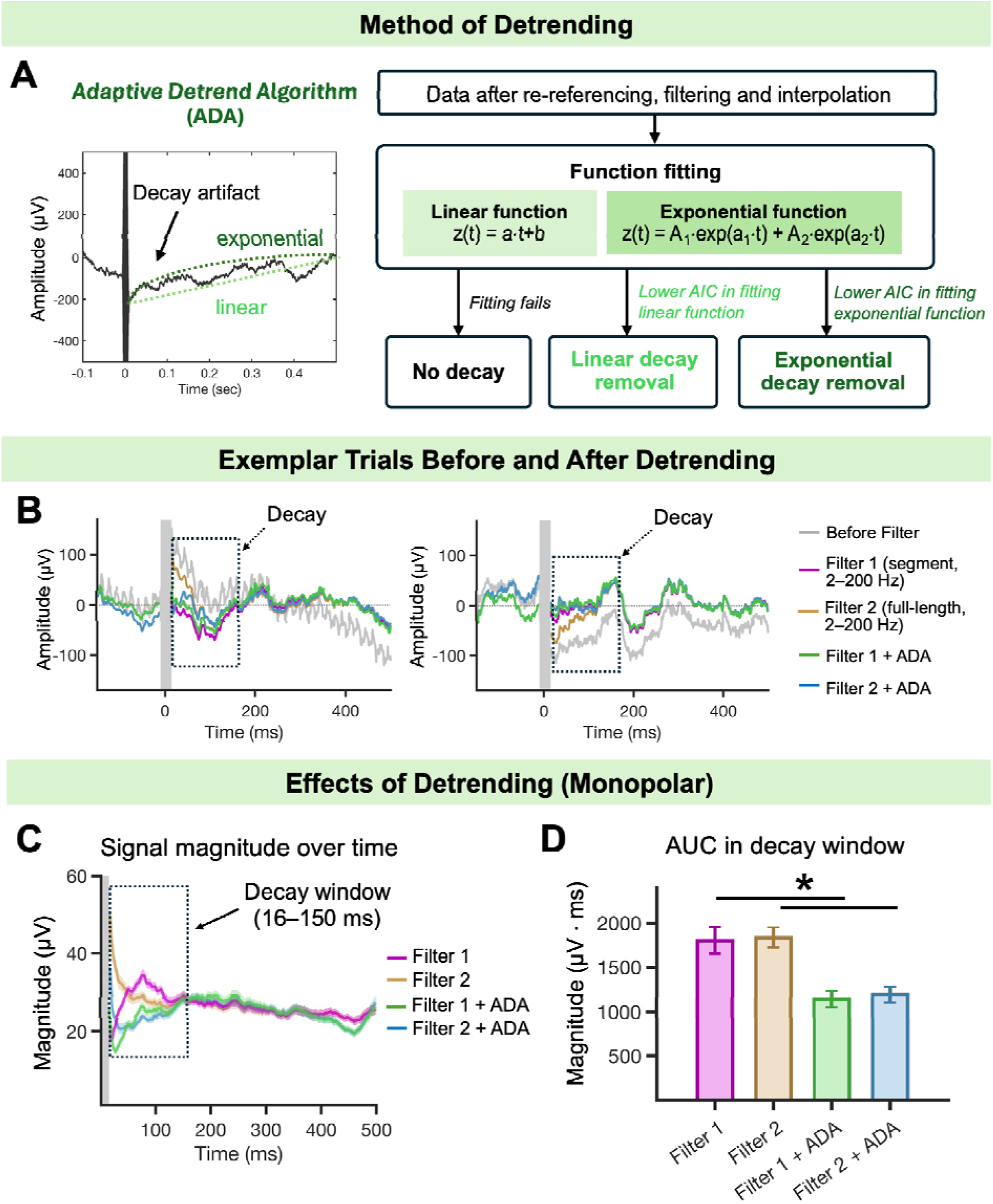
Method and Effects of Detrending. (A) Schematic of the adaptive detrending algorithm (ADA). (B) Example trials from two contacts across two participants. In both examples, ADA identified and removed potential decay artifacts within the early post-TMS window. (C) Trial-level averaged signal magnitudes before and after ADA processing on monopolar referenced data. Differences were primarily observed within the [16, 150 ms] time window (see Figure S5 for results in bipolar and common-average referenced data). (D) Contact-level statistics of area under the curve (AUC) within the decay window. Error bars represent the standard error across contacts. A total of 52 contacts were included in the statistical analysis, with three outlier contacts excluded for overwhelmingly large values (over three scaled mean absolute deviation from the group average). Statistically significant differences are indicated by bars (p_FDR_ < .05), confirming that ADA effectively attenuated post-TMS decay artifacts.

To quantify this effects, we computed the trial-level time-resolved signal magnitudes before and after ADA. Across all referencing and filtering conditions, differences were evident within the 16–150 ms time window (Figure 5C and Figure S5A), consistent with the typical decay duration reported in prior TMS-EEG studies (Hernandez-Pavon et al., 2022; Wu et al., 2018). Contact-level statistical analyses confirmed that ADA significantly reduced the AUC within this window across all conditions (ts < –3.17, p_FDR_ < .005; Figure 5D, S5). Besides, ADA after segment-based filtering yielded significantly lower AUC than after full-length filtering under common-average referencing condition (t(52) = –2.50, p_FDR_ = .023), but not under monopolar or bipolar referencing conditions (p_uncorrected_ > .31). Overall, these results support ADA as an effective and critical step for mitigating early post-TMS decay artifacts.

### 4. Effects of Full Preprocessing on Contact-Level iTEPs

After full preprocessing, only a small proportion of trials (<2%; see Table S2) were flagged as noisy and rejected based on the threshold criteria (see Methods). Contact-level iTEPs were then computed by averaging the remaining trials. Figure 6 presents iTEPs from three representative contacts across three participants, showing signals before and after preprocessing with different referencing and filtering methods.

**Figure 6.**
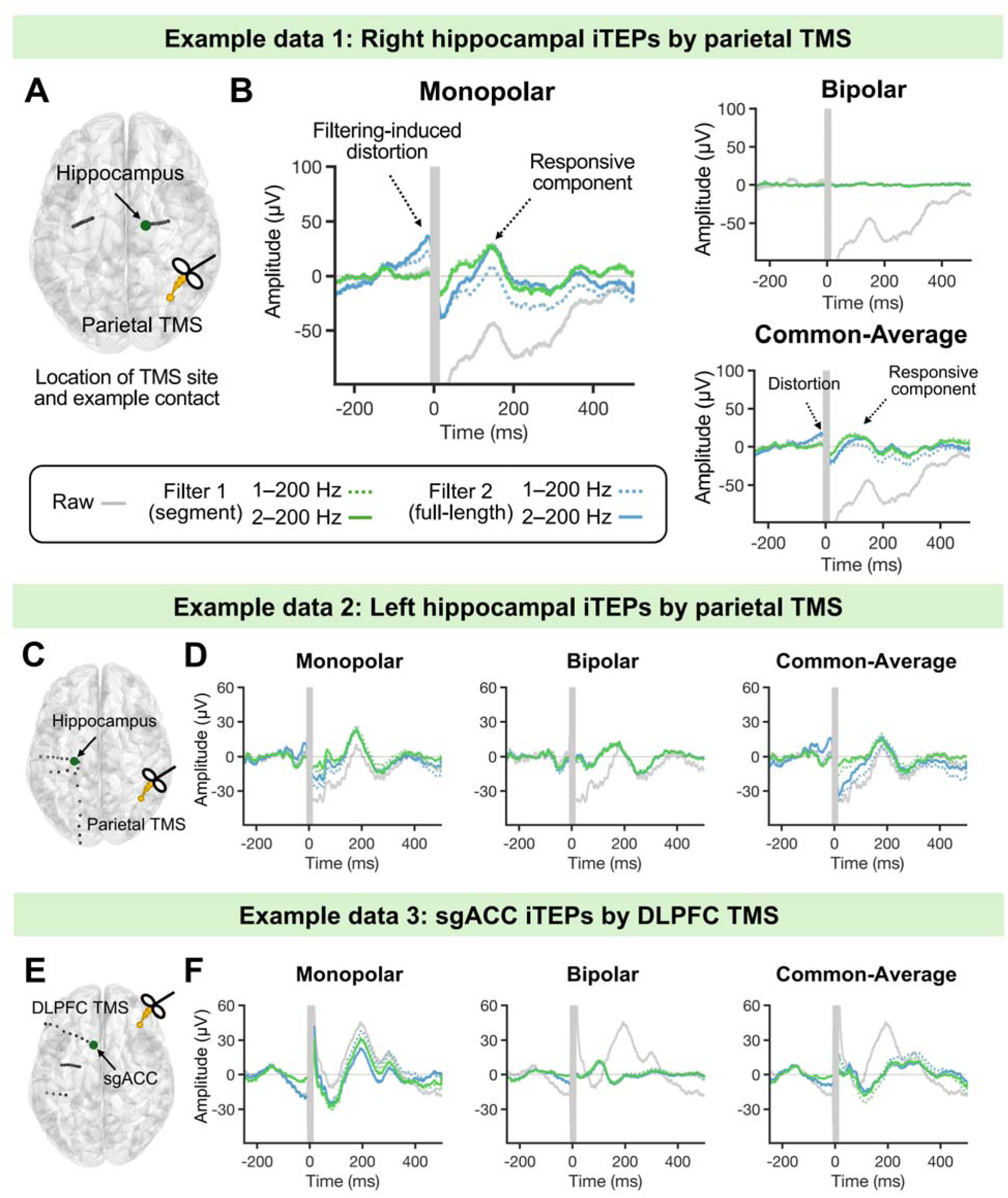
Contact-Level iTEPs Following Full Preprocessing Pipeline Across Methodological Choices. (A) Location of the TMS stimulation site and the exemplar hippocampus iEEG contact. (B) Hippocampal contact-level iTEPs before and after preprocessing using different referencing and filtering methods. Large artifacts present in the raw data were effectively attenuated after preprocessing. Notably, re-referencing substantially altered both waveform and amplitude: bipolar re-referencing reduced overall amplitude and suppressed prominent responses observed with monopolar or common-average methods. Segment-based filtering outperformed full-length filtering by better reducing residual post-TMS decay and minimizing pre-TMS distortions. The choice of bandpass setting (1–200 Hz vs. 2–200 Hz) had minimal impact with segment-based filtering but showed noticeable differences with full-length filtering. (C–F) Two additional exemplar iTEPs from two contacts from another two participants.

The first example (Figure 6A–B) shows hippocampal iTEPs elicited by parietal TMS. Raw data contained prominent artifacts lasting up to 500 ms, which were largely attenuated after preprocessing. However, the choice of preprocessing method notably affected iTEPs morphology and amplitude. In particular, re-referencing had a strong impact. The bipolar re-referencing reduced overall amplitude and attenuated a prominent response at ∼170 ms as observed with monopolar referencing (exceeding 3 pre-stimulation baseline SD). The common-average re-referencing preserved more signals but altered the waveform shape and shifted the peak earlier.

Differences between filtering strategies were most evident near the TMS artifact window. In both monopolar and common-average referencing conditions, full-length filtering introduced spurious artifacts during the pre-TMS interval (approximately –100 ms to –11 ms), whereas segment-based filtering better suppressed post-TMS decay and yielded signals that were closer to zero. The choice of bandpass setting (1–200 Hz vs. 2–200 Hz) had minimal impact when using segment-based filtering, but affected both pre-and post-TMS intervals under full-length filtering. These effects were less evident in bipolar referencing condition, likely due to overall weaker signals after bipolar subtraction. Similar findings were observed in the other two exemplar contacts (Figure 6C–F; see Figure S6–S8 for results across all contacts). Together, these findings highlight the effectiveness of our preprocessing pipeline and illustrate the influence of specific methodological choices, particularly the re-referencing and filtering strategies, on the resulting iTEPs.

### 5. Minimal Impact of Preprocessing Step Orders on iTEPs

To assess whether the sequence of the key preprocessing steps influences iTEPs, we compared our primary pipeline (Order 1: re-referencing → filtering → detrending) with two alternative sequences: Order 2: filtering → detrending → re-referencing; Order 3: detrending → re-referencing → filtering). The second order was motivated by designing the re-referencing as the last step for removing the shared residual noise or artifacts, whether inherited from the raw data or introduced during preprocessing. The third order was further motivated by enhancing the subsequent filtering by early removal of decay artifacts (Rogasch et al., 2017). In both alternative orders, resampling and epoching were performed after these three key steps. In this analysis, filtering was implemented using the segment-based strategy and with the 2– 200 Hz bandpass setting.

In a representative contact (Figure S9), iTEPs were highly similar across the three step orders at both trial and contact levels, with only negligible differences observed within the early post-TMS window. We then extended to all contacts (Figures S10–S12) and confirmed that contact-level iTEPs remained largely consistent regardless of step order. Some contacts exhibited small amplitude differences within the first 200 ms, typically with lower iTEP amplitudes when re-referencing was the first processing step (e.g., Figure S10, Pt 3–C16/17). Despite this, the major components in the majority of contacts were preserved across all preprocessing orders. Together, these results indicate that the order of processing steps can have a modest impact on contact-level iTEPs. When all critical preprocessing steps are included, the step order is unlikely to substantially alter estimates of the temporal dynamics of evoked responses or the binary conclusions about the regional responsiveness.

## Discussion

In this study, we presented a practical preprocessing pipeline for concurrent spTMS-iEEG data, incorporating several critical steps of re-referencing, filtering, artifact interpolation, and detrending. Using both real and simulated data, we systematically evaluated the effects of each preprocessing step and examined how various methodological choices/parameters and step orders affected iTEPs. Our results demonstrated that the pipeline effectively mitigated various artifacts and noises, yielding cleaner data suitable for subsequent analyses. To our knowledge, this is the first study to provide a detailed investigation into the preprocessing methodologies for concurrent TMS-iEEG data. Our work thus serves as an important first step toward establishing a general preprocessing framework and provides valuable guidance for future TMS-iEEG studies in determining appropriate workflows and specific methodological choices.

### 1. Referencing Choice Requires Careful Consideration

Our analysis of real spTMS-iEEG data demonstrated that referencing choice can substantially influence the waveform morphology and amplitude of iTEPs, potentially leading to divergent conclusions regarding the responsiveness of target regions and the temporal characteristics of evoked responses. Simulation further showed that referencing performance depends on specific signal conditions, underscoring the importance of evaluating actual recordings when selecting an appropriate approach. Specifically, when recording signals are not highly similar across contacts, suggesting that the reference electrode is not strongly contaminated by hardware artifacts or large TMS-evoked responses, monopolar referencing can reliably preserve and reflect true neural signals. In such cases, re-referencing may obscure meaningful activity or introduce non-target components. For example, bipolar re-referencing may reduce sensitivity after subtracting signals from nearby contacts that partially capture the same source. This issue was evident in a real example (Figure 6B), where bipolar re-referencing appeared to eliminate genuine signals shared across adjacent contact located within the same anatomical region, potentially leading to false negatives (Parish et al., 2023). Conversely, when recordings are dominated by shared non-target activity, monopolar referencing may yield false positives due to contamination from global noise. Under such conditions, re-referencing becomes increasingly necessary for isolating true evoked responses.

In our simulations, the common-average method performed poorly overall. It is likely due to the limited number of contacts (four contacts), which hindered accurate estimation of a shared noise profile. In contrast, real data analysis with broader contact coverage (often >100 contacts) showed that it might effectively remove global noise while preserving target responses, particularly compared to the bipolar method (e.g., Figure 6B). To better reflect the performance of common-average re-referencing under more realistic conditions, future simulation studies may employ more complex models, incorporating more latent sources and a larger number of contacts. Nonetheless, researchers should be cautious with this method, especially in datasets with sparse or unevenly distributed contacts (Mercier et al., 2022). In practice, selecting a subset of contacts by data-driven approaches (Huang et al., 2024) may help mitigate bias introduced by high-amplitude or noise-free channels, thereby enabling more accurate estimation and isolation of shared noise.

Taken together, we recommend that researchers carefully inspect raw TMS-iEEG data before applying any re-referencing method and make informed decisions based on the characteristics of the real dataset, anatomical targets, and research goals. When recorded signals are not highly similar across contacts, we advocate for greater consideration of monopolar referencing. In this case, to further assess the specificity of TMS-evoked responses, comparisons across anatomically adjacent yet functionally unrelated contacts or regions might be informative. When signals are highly similar across contacts, re-referencing becomes increasingly critical. Bipolar re-referencing or other local schemes (e.g., Laplacian re-referencing) is preferable when the goal is to detect local neural activities with high spatial specificity (Mercier et al., 2022), while common-average re-referencing may better balance the retention of broader, regionally distributed responses with the suppression of globally shared components (Huang et al., 2024).

### 2. Preprocessing Strategies for Strong TMS Artifacts

Raw spTMS-iEEG data contain high-amplitude, non-physiological artifacts surrounding the stimulation pulse, often exceeding the amplifier’s dynamic range (Figure S1). These segments are too noisy for any analysis and should be excluded. If continuity or stationarity is required in the subsequent processing, interpolation is further needed. While the artifact characteristics were generally consistent across trials (Figure S1), their duration and shape might vary across electrodes, shafts, and participants, necessitating manual inspection for defining an appropriate artifact exclusion window. In our dataset, the artifact period lasted approximately 10 milliseconds or less, leading us to define the exclusion window as [–10, +15 ms]. A narrower window may suffice when actual artifact duration is shorter, helping to preserve more early signals. On the contrary, a wider exclusion window (e.g., until +25 ms) might be necessary when artifacts persist longer, or when downstream analyses (e.g., spectral analysis) require temporal continuity or signal stationarity over longer intervals, as in previous studies (Li et al., 2025; Solomon et al., 2024).

To mitigate the influence of TMS artifacts on filtering, we recommend a segment-based filtering strategy, in which the artifact window is excluded and filters are applied only to the remaining signal segments. This approach outperformed the strategy by applying filters over full-length data after artifact interpolation, as the latter was found to introduce distortions that may bias estimates of TMS-evoked responses. While segment-based filtering may raise concerns about edge effects, i.e., distortions arising from the absence of data at the segment boundaries (Widmann and Schröger, 2012), our results show they can be effectively managed using standard padding techniques such as mirror padding (e.g., as implemented in filtfilt.m in MATLAB).

When designing the bandpass filter, we recommend setting the low cutoff frequency to 2 Hz, rather than a lower threshold such as 1 Hz. This helps reduce contamination from stimulation-related periodicity (0.5 Hz in this study) and better manage edge effects in short signal segments. In addition, analyses on real data showed that TMS-evoked responses observed with a 1-Hz filter were preserved using a 2-Hz filter (Figures S6–S8), alleviating concerns about the potential loss of meaningful signals. In this study, we used Butterworth IIR filters for both bandpass and notch filtering. Future studies may explore alternative filter designs, such as FIR filters, Chebyshev IIR filters, or adaptive filtering approaches, to further optimize filtering performance in spTMS-iEEG preprocessing.

### 3. Approaches for Removing Long-Lasting Decay Artifacts

The removal of decay artifacts, often persisting for over one hundred milliseconds, is essential for accurately characterizing iTEPs. In our pipeline, ADA was specifically implemented for this purpose and effectively identified and removed decay artifacts, particularly within 150 ms post-TMS window. Interestingly, bandpass filtering provided a complementary benefit in attenuating these post-TMS decay artifacts, especially when applied using the segment-based strategy. This was evidenced by more than 80% of trials showing reduced signal magnitude at the offset of the TMS artifact, along with a substantial reduction in contact-level signal magnitude (approximately one-fourth of the level before filtering). This effect is likely attributable to the temporal profile of decay artifacts, which resemble low-frequency drifts characterized by a sharp deflection at the TMS offset that gradually returns to baseline over time. Such low fluctuations are attenuated when falling below the lower cutoff frequency of the bandpass filter (2 Hz). Together, segment-based filtering and ADA provided a powerful combination for mitigating decay artifacts, enable more accurate characterization of evoked responses.

Considering the high amplitude of decay artifacts near the TMS artifact, we further explored whether extending the artifact exclusion window could further improve preprocessing. We compared the iTEPs obtained using a 15-ms vs. 25-ms exclusion window. As shown in example data (Figure S13A–B), the 15-ms window retained more early signals (16–25 ms), but iTEPs beyond this range were highly similar at both the trial and contact levels. Analyses across all contacts (Figures S14–S16) confirmed minimal differences beyond 25 ms and near-identical signals beyond 50 ms. These results suggest that our pipeline effectively mitigates decay artifacts and is robust to moderate variations in the artifact exclusion window.

Although the 15-ms window preserves more early signal within 16–25 ms, caution is warranted when interpreting these early responses. For example, some contacts (e.g., Figure S14, Pt 3–C6) exhibited large early waveforms that peaked at 16 ms and then decayed monotonically. This morphology more closely resembled decay artifacts in raw data (see Figure S10) than canonical early neural responses such as the N1 component typically observed following intracranial electrical stimulation (Keller et al., 2014). Therefore, it may be inappropriate to directly attribute these signals to genuine TMS-evoked activity. Given the limited understanding of the early TMS-evoked responses in distant brain regions, and the generally high noise amplitude within this early time window, further empirical and simulation work is needed to refine both interpretation and preprocessing strategies.

### 4. Conclusion and Future Direction

Concurrent spTMS-iEEG has emerged as a powerful tool for probing causal brain connectivity, yet its preprocessing poses significant challenges due to substantial TMS-related artifacts. In this study, we presented a practical preprocessing pipeline incorporating several key steps of re-referencing, filtering, interpolation and detrending. We demonstrated that this pipeline effectively attenuates various artifacts and noise, providing a solid foundation for subsequent iTEP analyses. Using both real and simulated datasets, we systematically evaluated the effects of each step and compared alternative methodological choices. Our findings particularly highlight the critical influence of referencing methods and filtering strategies on iTEP outcomes, offering practical guidance for selecting appropriate preprocessing approaches in future studies. Overall, this work represents an important step toward establishing a principled preprocessing framework. We hope it encourages broader attention to and methodological development in TMS-iEEG research, ultimately advancing our understanding of brain organization and TMS mechanisms.

Future work could further validate and extend our work along several lines. First, the generalizability of this pipeline should be assessed across larger and more diverse datasets, which may differ in experimental protocols (e.g., TMS intensity and frequency), recording environments, and amplifier systems. Second, it is important to systematically validate the pipeline’s applicability to ECoG-based iEEG data, as this study mainly focused on sEEG. Differences between these modalities, such as electrode material, anatomical coverage, and orientation relative to the neural tissue, may affect both signal and noise characteristics (Miller et al., 2023). While preliminary results from two exemplar ECoG contacts (Figure S17) support the utility of this pipeline, broader validation and cross-modality comparisons are warranted. Third, future studies could investigate and incorporate advanced data-driven techniques, such as ICA-or PCA-based approaches for re-referencing or artifact removal (Alexander et al., 2019; Michelmann et al., 2018). These methods would expand the available methodological toolkit and improve the flexibility and scalability of TMS-iEEG preprocessing across diverse datasets.

## Funding

JJ is supported by the National Institutes of Health (R01MH136197) and the Brain and Behavior Research Foundation Young Investigator grant (29441). CJK is supported by grants from the National Institutes of Health (R01MH126639, R01MH129018), and Burroughs Welcome Fund Career Award for Medical Scientists. NTT is supported by grants from the National Institute of Mental Health (1K23MH125145), and the Brain and Behavior Research Foundation (31275 and 32270). ADB is supported by the National Institutes of Health (R21MH120441, R01NS114405) and Roy J. Carver Trust. Other funding includes NIMH R01MH132074 and R01MH139650 (CJK, NTT, ADB). This work was conducted, in part, on an MRI instrument funded by 1S10OD025025-01.

## CRediT authorship contribution statement

Zhuoran Li: Writing – review & editing, Writing – original draft, Software, Methodology, Formal analysis, Data curation, Conceptualization.

Xianqing Liu: Writing – review & editing, Writing – original draft, Methodology, Data curation, Conceptualization.

Joshua Tatz: Writing – review & editing, Writing – original draft, Methodology, Conceptualization

Umair Hassan: Writing – review & editing, Software, Methodology, Conceptualization

Jeffrey B Wang: Writing – review & editing, Software, Methodology, Conceptualization Corey J. Keller: Writing – review & editing, Resources, Methodology, Funding acquisition, Conceptualization

Nicholas T. Trapp: Writing – review & editing, Resources, Investigation, Methodology, Funding acquisition, Conceptualization

Aaron D. Boes: Writing – review & editing, Writing – original draft, Supervision, Resources, Project administration, Investigation, Funding acquisition, Conceptualization Jing Jiang: Writing – review & editing, Writing – original draft, Supervision, Resources, Project administration, Methodology, Funding acquisition, Conceptualization

## Declaration of competing interest

C.J.K. holds equity in Alto Neuroscience, Inc, and is a consultant for Flow Neuroscience.

## Acknowledgement

We would like to acknowledge the patients and families who graciously agreed to participate in this research, as well as colleagues who assisted with data collection and provided feedback on the project, including Joel Bruss, Ben Pace, Brandt Uitermarkt, Ariane Rhone, Haiming Chen, Kirill Nourski, Joel Berger, Hiroyuki Oya, and Chris Garcia.

