## Supplementary information for "A Practical Preprocessing Pipeline for Concurrent TMS-iEEG: Critical Steps and Methodological Considerations"

### Supplementary Materials

#### Supplementary Methods

##### Real Dataset

We analyzed data from three neurosurgical patients (2 female, ages 23 and 24; 1 male, age 34). These patients with medically intractable epilepsy were admitted to the University of Iowa Hospitals and Clinics for monitoring with intracranial electrode implantation to localize their seizure focus. All experimental procedures were approved by the University of Iowa Institutional Review Board (IRB) and were conducted in accordance with ethical principles in the Declaration of Helsinki. Written informed consent was obtained from all patients.

During the concurrent spTMS-iEEG experiment, 50 single pulses of TMS were delivered at a frequency of 0.5 Hz. Among the three patients, two received stimulation at the right parietal cortex, and one at the right dorsolateral prefrontal cortex (DLPFC). Parietal TMS sites were individually defined based on the parietal location showing maximal resting-state functional connectivity (FC) to the hippocampus [1]. The DLPFC target was near the Beam F4 location, chosen as the right-hemisphere homologue of a previously established left DLPFC coordinate [2]. Stimulation was delivered at 120% of motor threshold (MT), or 100% MT if 120% was not tolerated. During the experiment, Brainsight Neuronavigation System (Rogue Research, Montreal, Quebec, Cannada) and the BEST toolbox [3] were used for neuronavigation and TMS protocol control, respectively.

iEEG data were simultaneously recorded using a multichannel data acquisition system (ATLAS, Neuralynx, Tucson, AZ), with a reference electrode placed in patients' subgaleal space. Signals were sampled at 8 kHz or 4 kHz and filtered online with 1–2000 Hz acquisition filters (−6 dB, 256 tap length). Depth electrodes (Stereo-electroencephalography, sEEG) and/or subdural grid/strip arrays (electrocorticography, ECoG) (Ad-Tech Medical, Racine, WI) were implanted in all three patients. We specifically selected sEEG shafts covering the deep brain regions of interest (ROIs), including the hippocampus, amygdala, and sgACC. Anatomical labels for each sEEG contacts were first determined using MNI-space coordinates with reference to established atlas or mask. Specifically, the Harvard-Oxford Structural Atlas at a 25% threshold in FSL [4] was used to identify contacts located in the hippocampus or amygdala. A sgACC mask generated in our previous study [5] was used to identify contacts located in the sgACC. All identified contacts were then manually verified in native space. Subsequently, all other contacts within the same shaft and inside the brain were also included. Additional methodological details, including implantation imaging, electrode coordinate computation, etc., are consistent with and described in our prior studies [5, 6]. Based on this selection strategy, we identified 55 sEEG contacts across three participants. In this study, we systematically assessed the effects of each preprocessing step and examine how different preprocessing choices affect the resulting iTEPs.

**Simulated Data**

The stimulation procedures were adapted from a previously published study (Michelmann et al., 2018). First, we defined three latent sources by using real spTMS-iEEG data from three different patients. It allowed us to preserve the temporal-spectral characteristic of biological data while avoiding data dependencies. For each source, a 1-sec segment was selected from a random contact between 16–1015 ms after a random TMS pulse. This time window was chosen to ensure the absence of strong TMS artifacts. These three data sources served as source 1 (S1), source 2 (S2) and reference source (R). They were all demeaned and scaled to unit variance, with the reference source scaled to 10% of the strength of the other two sources.

Then, we defined a linear mixture model to simulate the data recorded in a shaft (C1-C4). S1 and S2 were placed on contacts C1 and C4 respectively. They could spread into the neighboring contacts with varying strengths of $1/a$, $1/a^2$, and $1/a^3$ ($a > 1$), reflecting a decay spatial spreading of potential with distance. The parameter a was decreased in 40 logarithmic steps from 10 to 1.02. A smaller a, i.e., a higher level of $1/a^3$, means the effects spread to a greater extent from the second source. Additionally, each signal was referenced against the reference signal with a constant coefficient equaling –1. Gaussian pink noise was further added to each contact, with its variance varied across 40 linearly spaced levels (0–100% of the source signal variance), to simulate different noise conditions. Noise was generated separately for each contact to ensure independence.

Next, we applied three typical referencing methods to the simulated data, including monopolar referencing (raw), bipolar re-referencing (subtraction of adjacent contact), and common-average re-referencing (subtraction of the average signal across four contacts). For the bipolar re-referencing, three new virtual contacts were created: C1' = C1 – C2, C2' = C2 – C3, C3' = C3 – C4.

Last, we computed the sensitivity and specificity for each referencing method. Sensitivity was defined as the mean Pearson correlation between the referenced signal at the two ends of the shaft (i.e., C1/C4 for monopolar and common-average methods; C1′/C3′ for bipolar method) and its nearby true source (i.e., S1/S2). Specificity was defined as 1 minus the mean Pearson correlation with the opposite, distant source (i.e., S1-C4, S2-C1 for monopolar and common-average methods; S1-C3′, S2-C1′ for bipolar method). To obtain a robust estimate, we repeated the simulation 1000 times for each level of a and each level of noise.

**Table S1. Demographic and Experimental Information of Patients**

| Patient | Age | Sex | Handed ness | Ethnicity | Education (yr) | Age of onset (yr) | Onset of Epilepsy | Lesions | MT (% machine output) | Stimulation Intensity (% MT) | # Contacts | Implantation Type |
|---|---|---|---|---|---|---|---|---|---|---|---|---|
| Pt 1 | 34 | M | - | non-Hispanic White | 12 | 20 | left occipital | focal cortical dysplasia type IIb | 59% | 100% | 192 | ECoG & sEEG |
| Pt 2 | 23 | F | R | non-Hispanic White | 12 | 16 | left lateral posterior parietal | none | 59% | 120% | 148 | ECoG & sEEG |
| Pt 3 | 24 | F | R | non-Hispanic White | 14 | 19 | left hemispher e | none | 72% | 100% | 146 | sEEG |

**Table S2. Proportion of Noisy Trials Identified Across Preprocessing Choices**

Values represent the mean (standard deviation) of the proportion of noisy trials across 52 contacts under different preprocessing conditions. Here, three outlier contacts with >50% noisy trials in both raw and preprocessed data (Pt 2–C6; Pt 3–C18/C19) were excluded from analysis. In the raw data, due to the presence of large TMS artifacts, 100% of trials were classified as noisy across all contacts. When the artifact window was excluded, the average proportion of noisy trials dropped to 6.04%. Preprocessing further reduced this proportion across all methodological choices ($p_{FDR} < .001$), with the proportion falling below 2%.

| | Filter 1 (segment) | | Filter 2 (full-length) | | Raw |
|---|---|---|---|---|---|
| | 1-200Hz | 2-200Hz | 1-200Hz | 2-200Hz | |
| **Monopolar** | 1.50(3.38) | 0.69(1.77) | 1.46(3.63) | 1.08(2.76) | 6.04(7.50) |
| **Bipolar** | 1.66(8.01) | 1.28(6.51) | 1.49(7.74) | 1.23(6.78) | |
| **Common-Average** | 0.88(2.84) | 0.50(1.67) | 1.27(2.69) | 0.65(1.76) | |

**Figure S1. Strong TMS Artifacts Surrounding the Stimulation Pulse**

Across all contacts, strong artifacts lasted approximately 10 milliseconds or less. While their waveforms consistently exceeded the amplifier's dynamic range and were highly similar across trials, they varied across contacts, shafts, and participants. Based on these observations, the artifact window was defined as [–10, +15 ms].

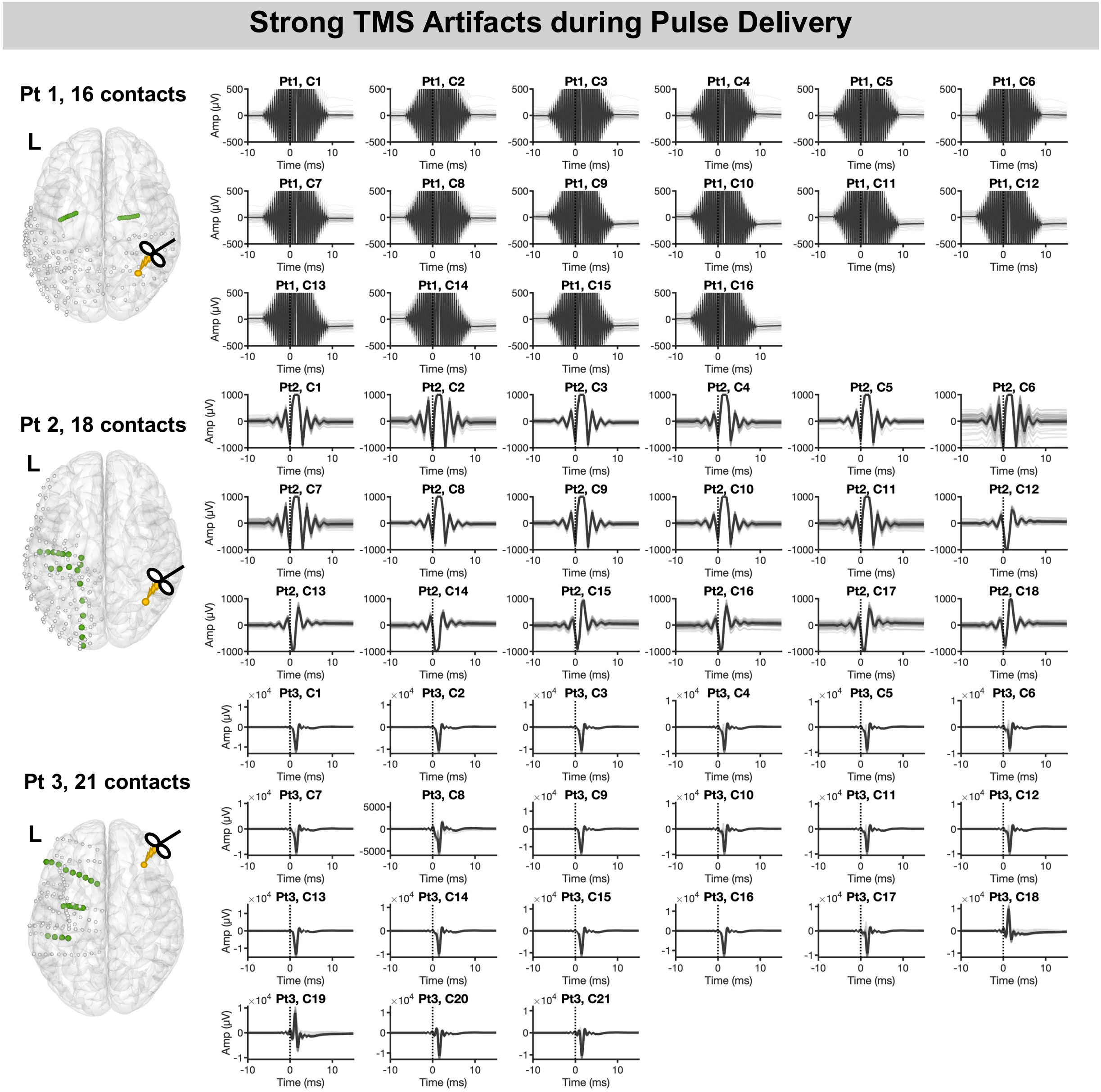

**Figure S2. Difference of Performance across Referencing Methods**

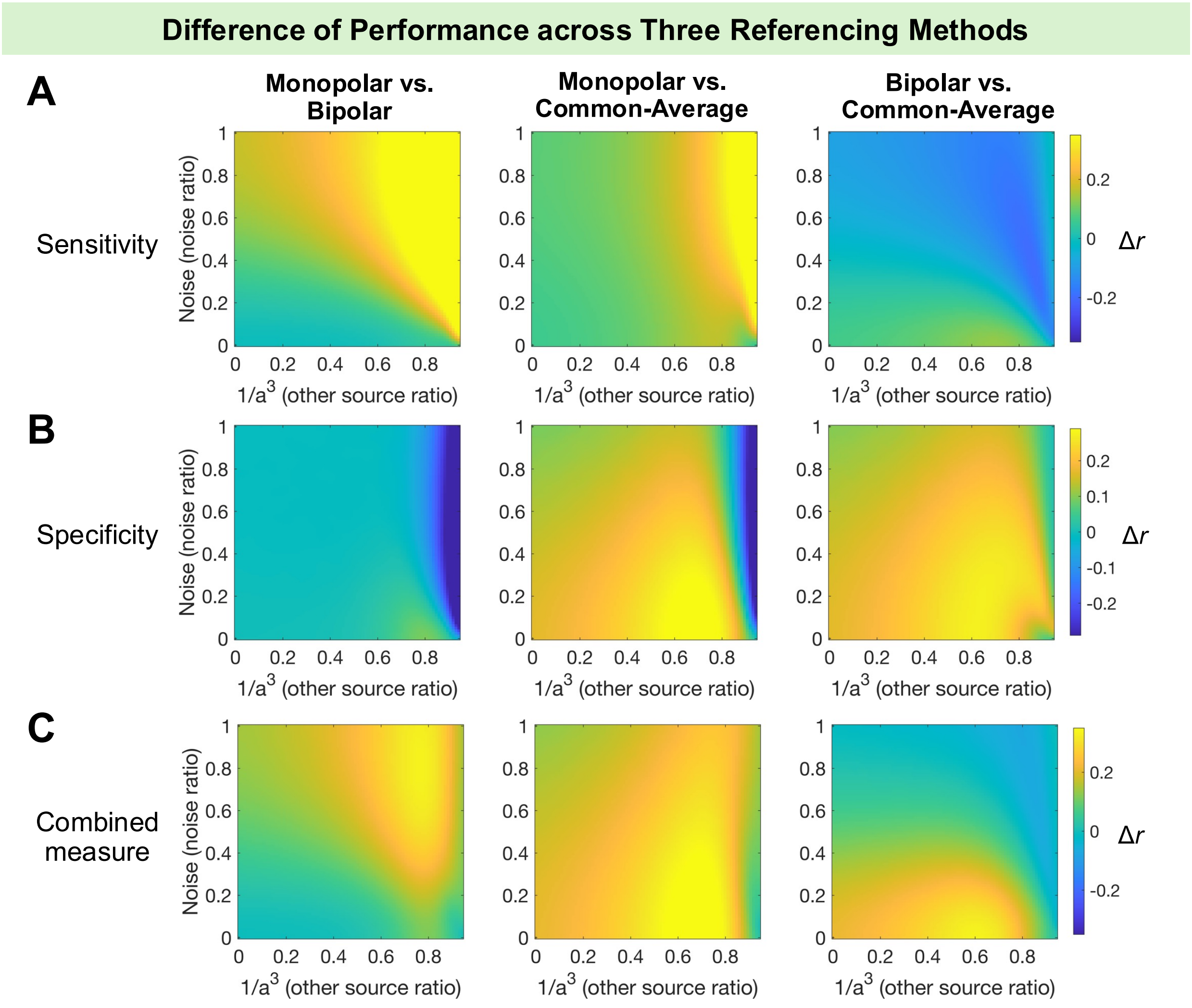

**Figure S3. Performance of Referencing Methods at Different Levels of Reference Source.**

The performance of the monopolar method declined as the reference source amplitude (variance) increased from 20% to 100% of the target source, based on results from N = 100 simulations.

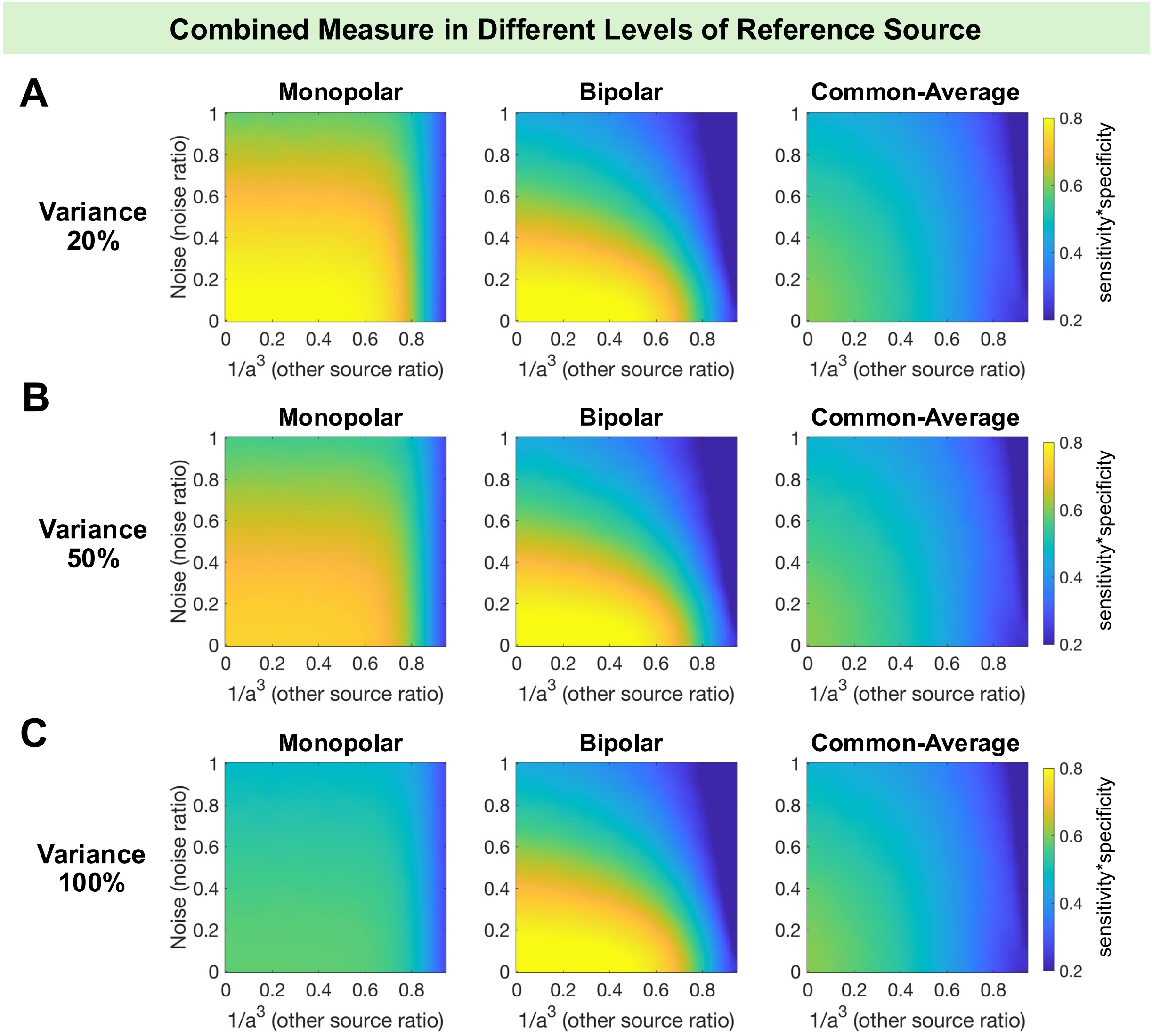

**Figure S4. Effects of Filtering and Interpolation on Bipolar and Common-Average Re-Referenced data**

For both bipolar and common-average re-referenced data, trial- and contact-level analyses demonstrated that segment-based filtering (Filter 1) more effectively attenuated artifacts immediately beyond the TMS artifact window while introducing minimal edge distortion, compared to full-length filtering (Filter 2). Results from 1-Hz and 2-Hz low cutoffs were generally similar, though the 2-Hz filter showed overall better performance. Specifically, in bipolar re-referenced data, the 2-Hz filter yielded lower signal magnitude at the TMS offset with segment-based filtering, with no significant differences in other conditions. In common-average re-referenced data, the 2-Hz filter resulted in lower magnitudes at both TMS onset and offset with segment-based filtering, and lower offset but higher onset magnitude with full-length filtering. Statistically significant contact-level differences are indicated by bars ($p_{FDR} < .05$).

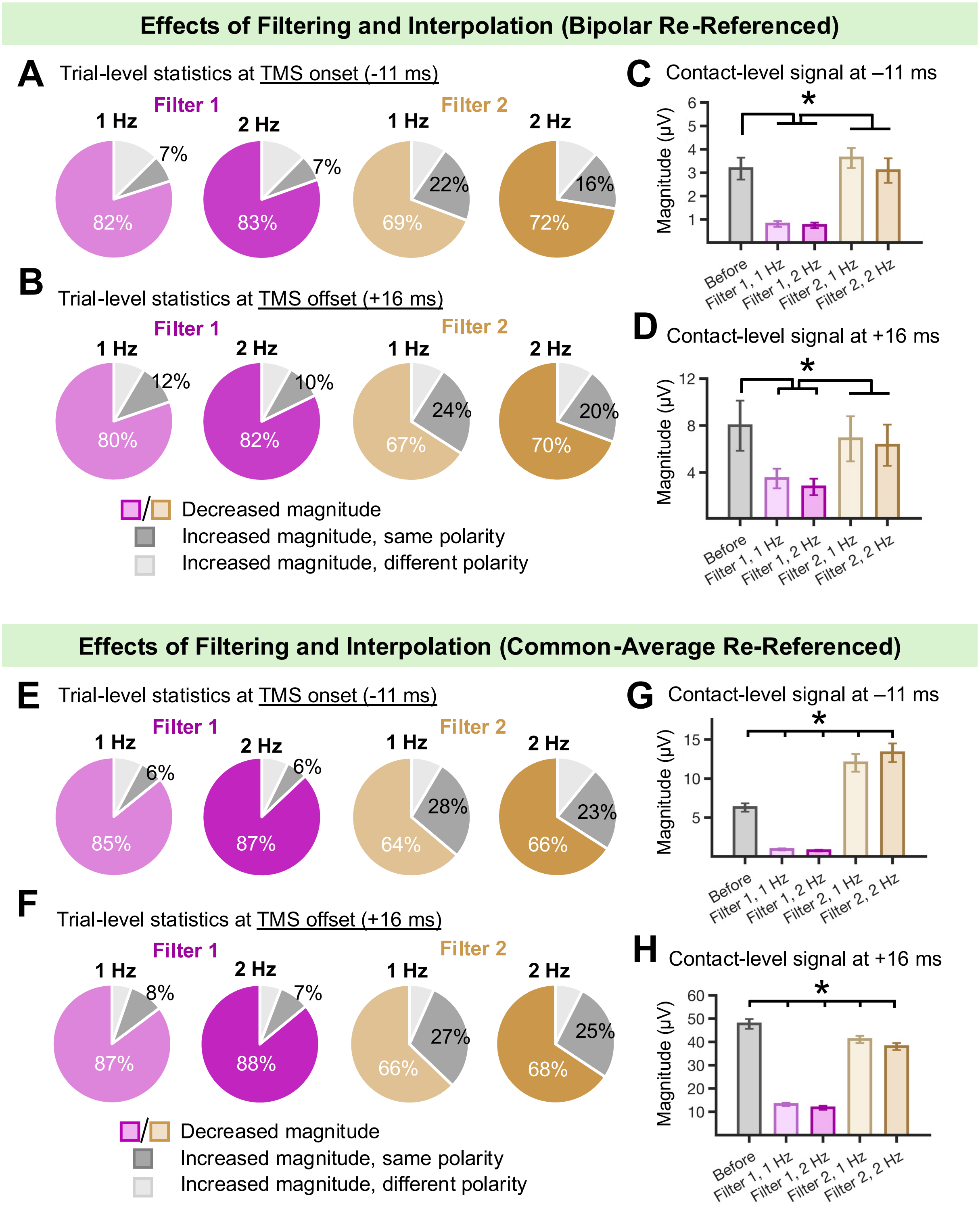

Effects of Filtering and Interpolation (Bipolar Re-Referenced)
A Trial-level statistics at TMS onset (-11 ms)
Filter 1
Filter 2
1 Hz
2 Hz
1 Hz
2 Hz
7%
82%
7%
83%
22%
69%
16%
72%
B Trial-level statistics at TMS offset (+16 ms)
Filter 1
Filter 2
1 Hz
2 Hz
1 Hz
2 Hz
12%
80%
10%
82%
24%
67%
20%
70%
Decreased magnitude
Increased magnitude, same polarity
Increased magnitude, different polarity
C Contact-level signal at –11 ms
Magnitude (µV)
*
Before
Filter 1, 1 Hz
Filter 1, 2 Hz
Filter 2, 1 Hz
Filter 2, 2 Hz
D Contact-level signal at +16 ms
Magnitude (µV)
*
Before
Filter 1, 1 Hz
Filter 1, 2 Hz
Filter 2, 1 Hz
Filter 2, 2 Hz
Effects of Filtering and Interpolation (Common-Average Re-Referenced)
E Trial-level statistics at TMS onset (-11 ms)
Filter 1
Filter 2
1 Hz
2 Hz
1 Hz
2 Hz
6%
85%
6%
87%
28%
64%
23%
66%
F Trial-level statistics at TMS offset (+16 ms)
Filter 1
Filter 2
1 Hz
2 Hz
1 Hz
2 Hz
8%
87%
7%
88%
27%
66%
25%
68%
Decreased magnitude
Increased magnitude, same polarity
Increased magnitude, different polarity
G Contact-level signal at –11 ms
Magnitude (µV)
*
Before
Filter 1, 1 Hz
Filter 1, 2 Hz
Filter 2, 1 Hz
Filter 2, 2 Hz
H Contact-level signal at +16 ms
Magnitude (µV)
*
Before
Filter 1, 1 Hz
Filter 1, 2 Hz
Filter 2, 1 Hz
Filter 2, 2 Hz

**Figure S5. Removal of Decay Artifacts by ADA**

Across all preprocessing conditions—regardless of referencing or filtering choices—ADA effectively identified and attenuated decay artifacts within the 16–150 ms post-TMS window, resulting in reduced area under the curve (AUC). Statistically significant differences are indicated by bars ($p_{FDR} < .05$).

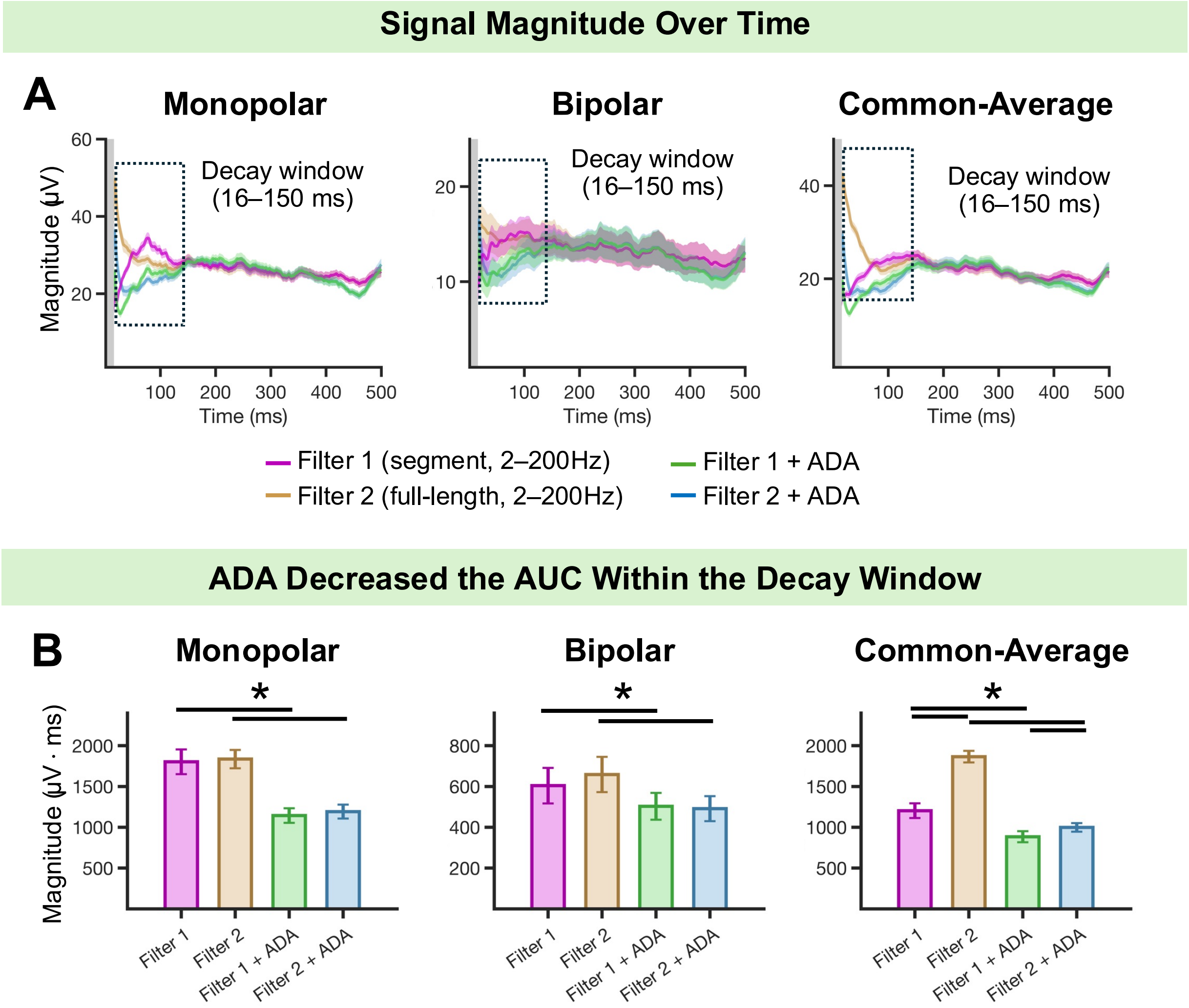

**Figure S6. Contact-level iTEPs Before and After Preprocessing (Monopolar Referencing)**

Compared to raw data, our preprocessing pipeline effectively mitigated artifacts and noise, yielding cleaner iTEPs. The iTEPs derived from different filtering choices were similar. Some contacts (e.g., Pt 2-C6, Pt 3-C18/19) remained noisy after preprocessing due to inherently noisy raw signals.

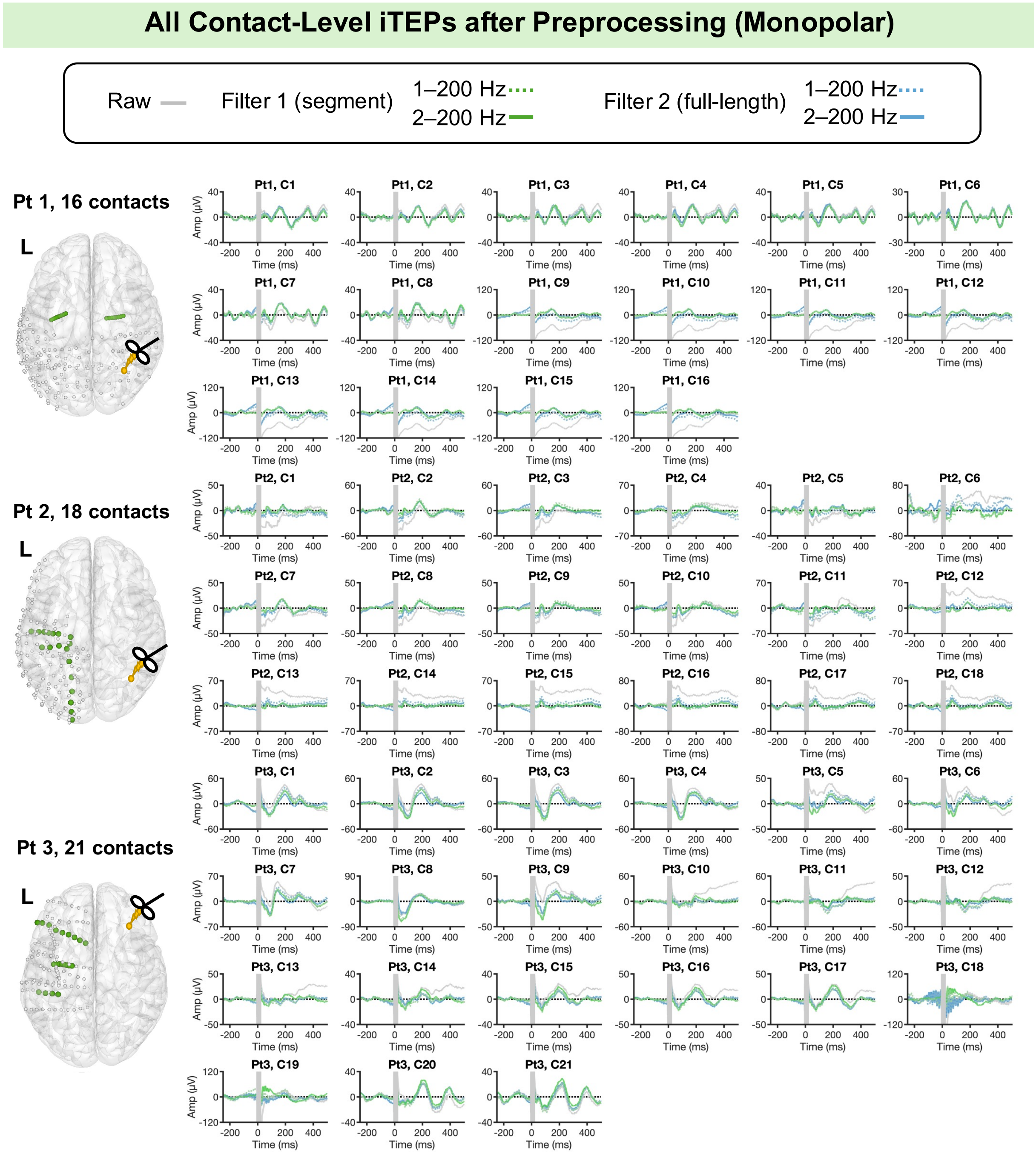

**Figure S7. Contact-level iTEPs Before and After Preprocessing (Bipolar Re-Referencing)**

Bipolar re-referencing substantially reduced signal amplitude and altered waveform morphology. The iTEPs derived from different filtering choices were similar.

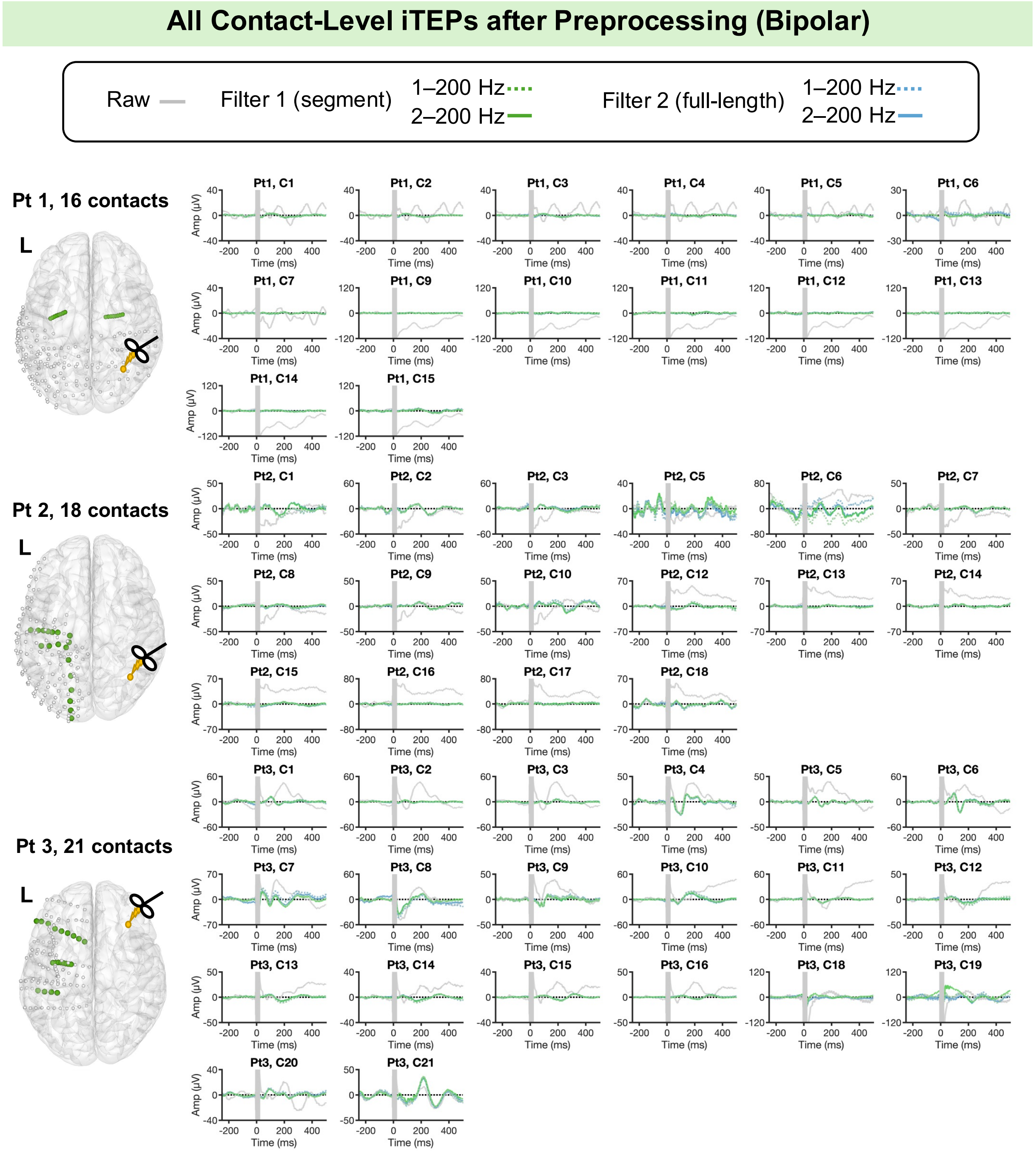

**Figure S8. Contact-level iTEPs Before and After Preprocessing (Common-Average Re-Referencing)**

Common-average re-referencing changed waveform morphology but can preserve some components observed in the raw data. The iTEPs derived from different filtering choices were similar.

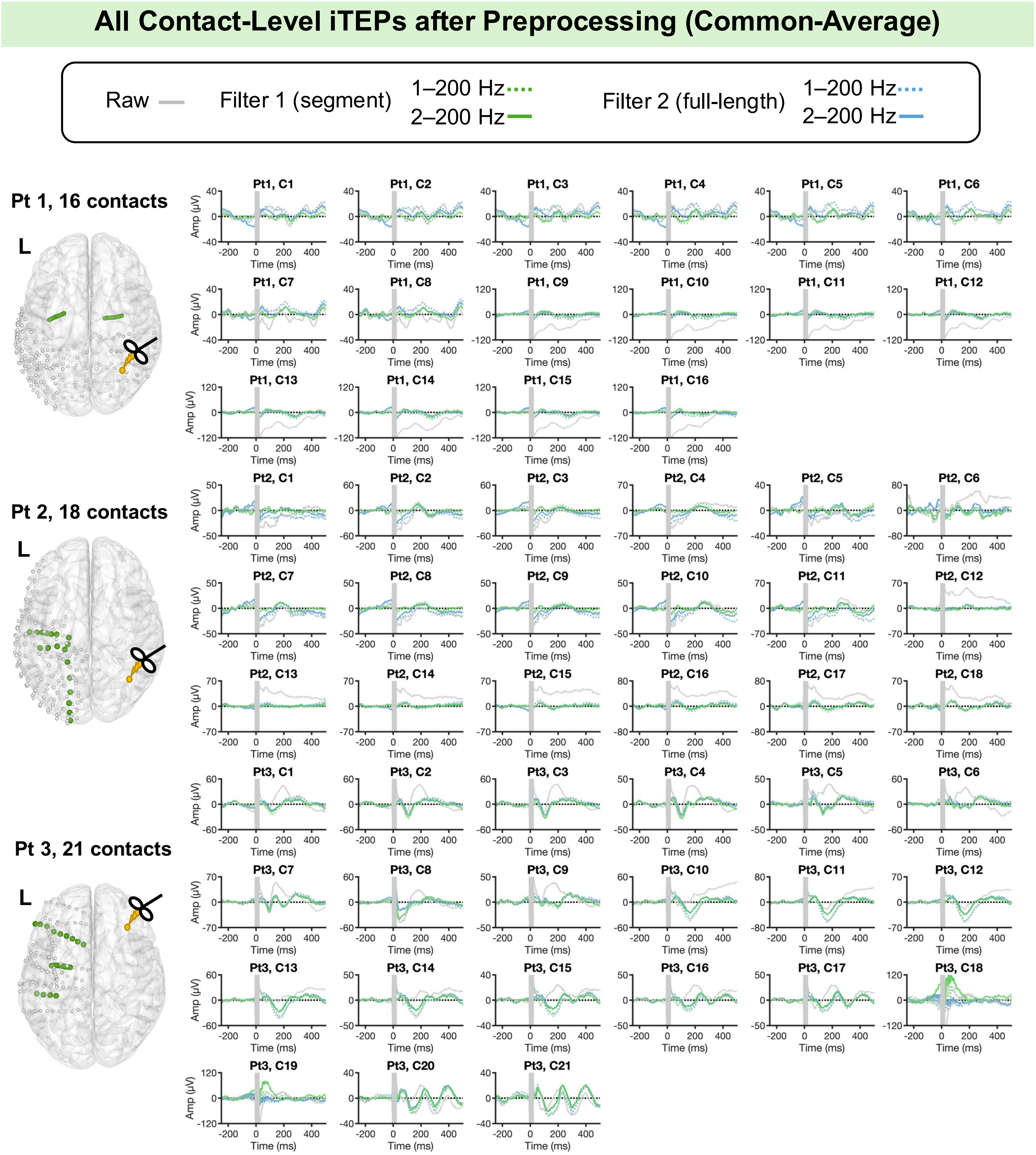

**Figure S9. Examples Illustrating the Effects of Preprocessing Step Order on Trial-Level and Contact-Level iTEPs**

Three preprocessing pipelines with different step orders were evaluated and compared: (1) re-referencing → filtering → detrending (primary pipeline); (2) filtering → detrending → re-referencing; (3) detrending → re-referencing → filtering. Here, filtering was achieved by segment-based strategy, with a 2–200 Hz as the bandpass filtering range. The resulting iTEPs were highly similar across all orders at both the trial and contact levels.

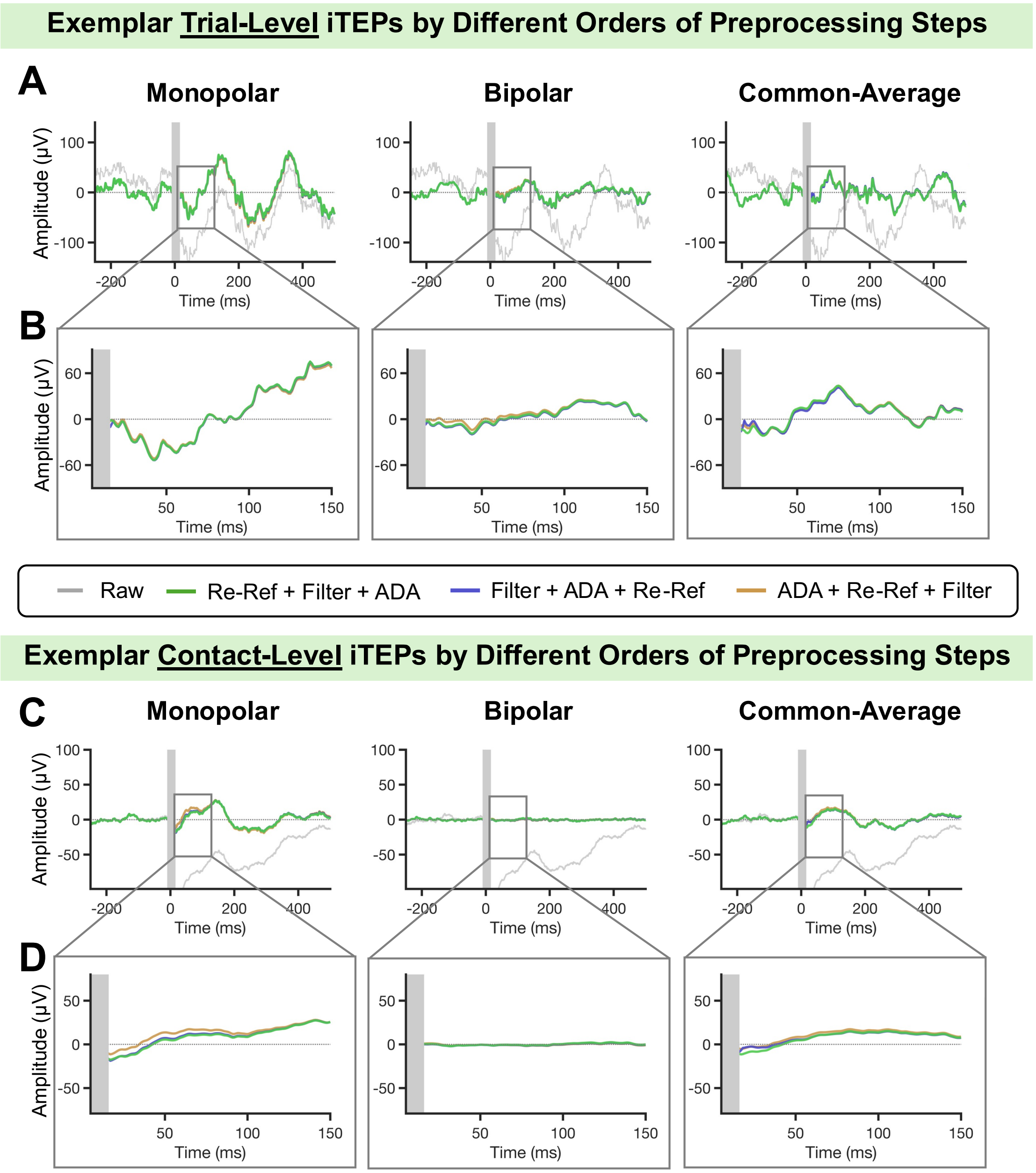

**Figure S10. Contact-level iTEPs by Different Preprocessing Orders (Monopolar Referencing)**

Compared to raw data, three preprocessing pipelines effectively mitigated artifacts and noise, yielding cleaner iTEPs. While modest differences were observed within the first 200 ms in some contacts (e.g., from Pt 3), the iTEPs derived from the three preprocessing orders are highly similar.

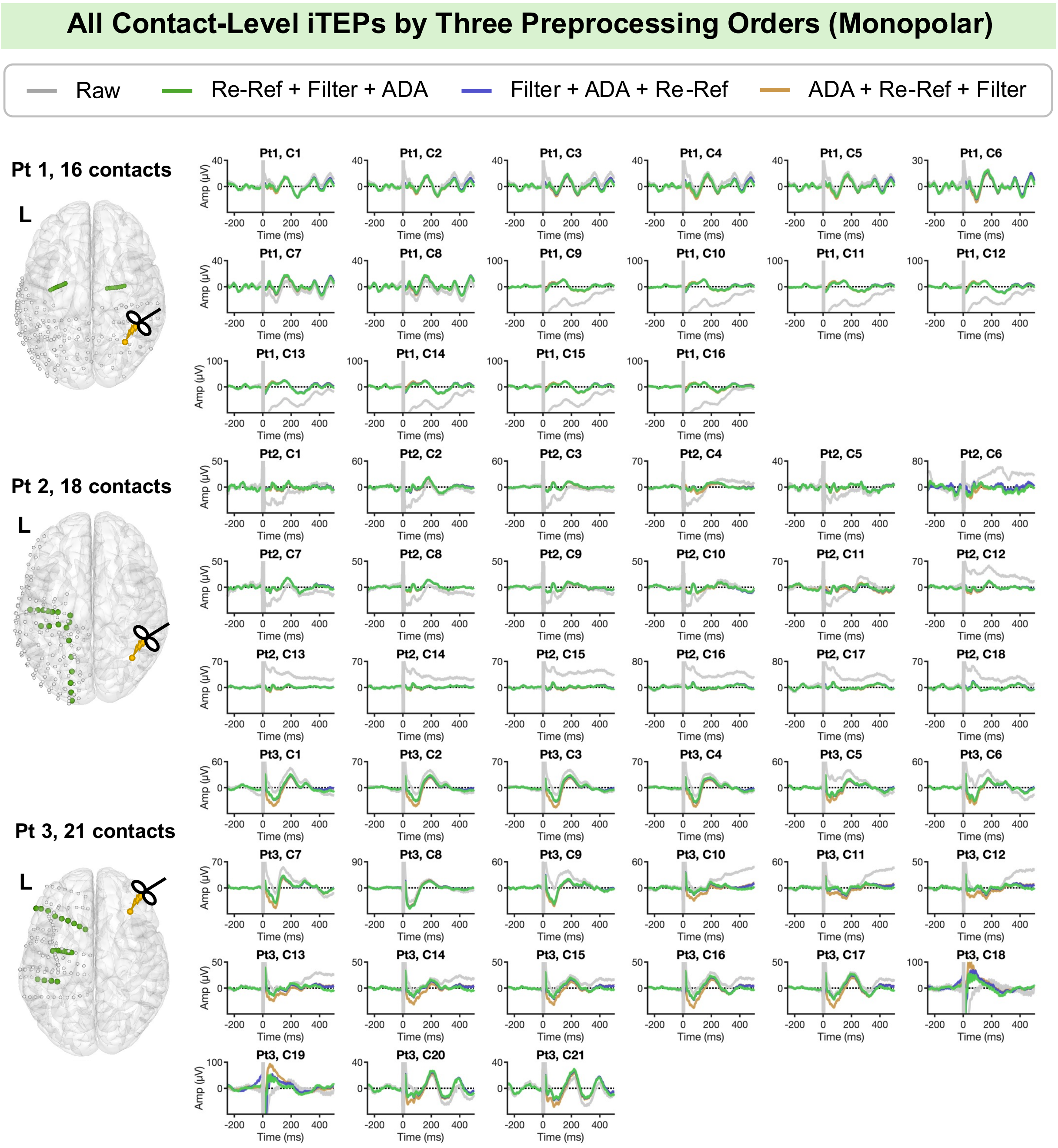

**Figure S11. Contact-level iTEPs by Different Preprocessing Orders (Bipolar Re-Referencing)**

The iTEPs derived from the three preprocessing orders are highly similar.

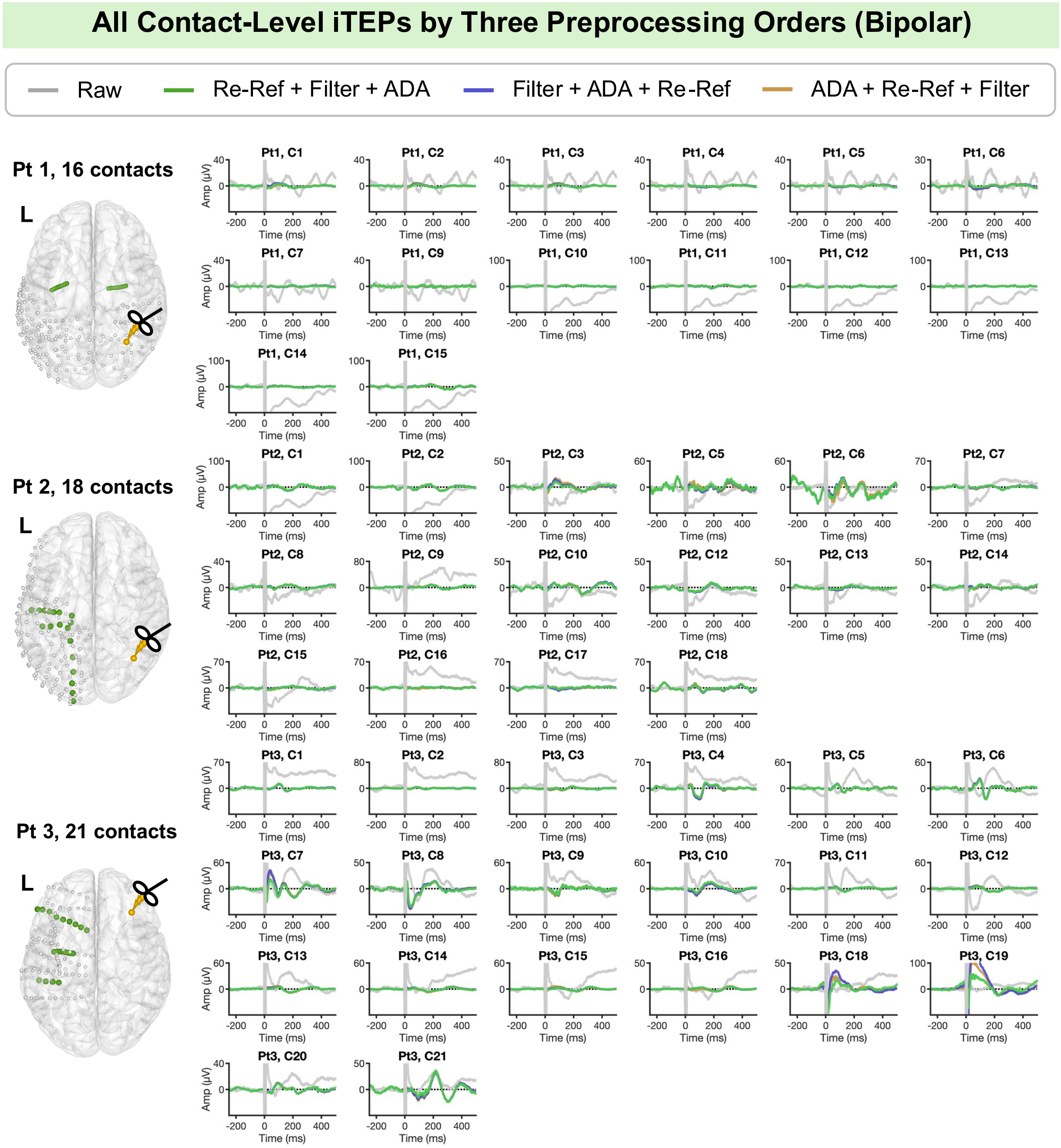

**Figure S12. Contact-level iTEPs by Different Preprocessing Orders (Common-Average Re-Referencing)**

iTEPs derived from the three preprocessing orders are highly similar, with minor differences were observed within ~100 ms in some contacts (e.g., from Pt 3).

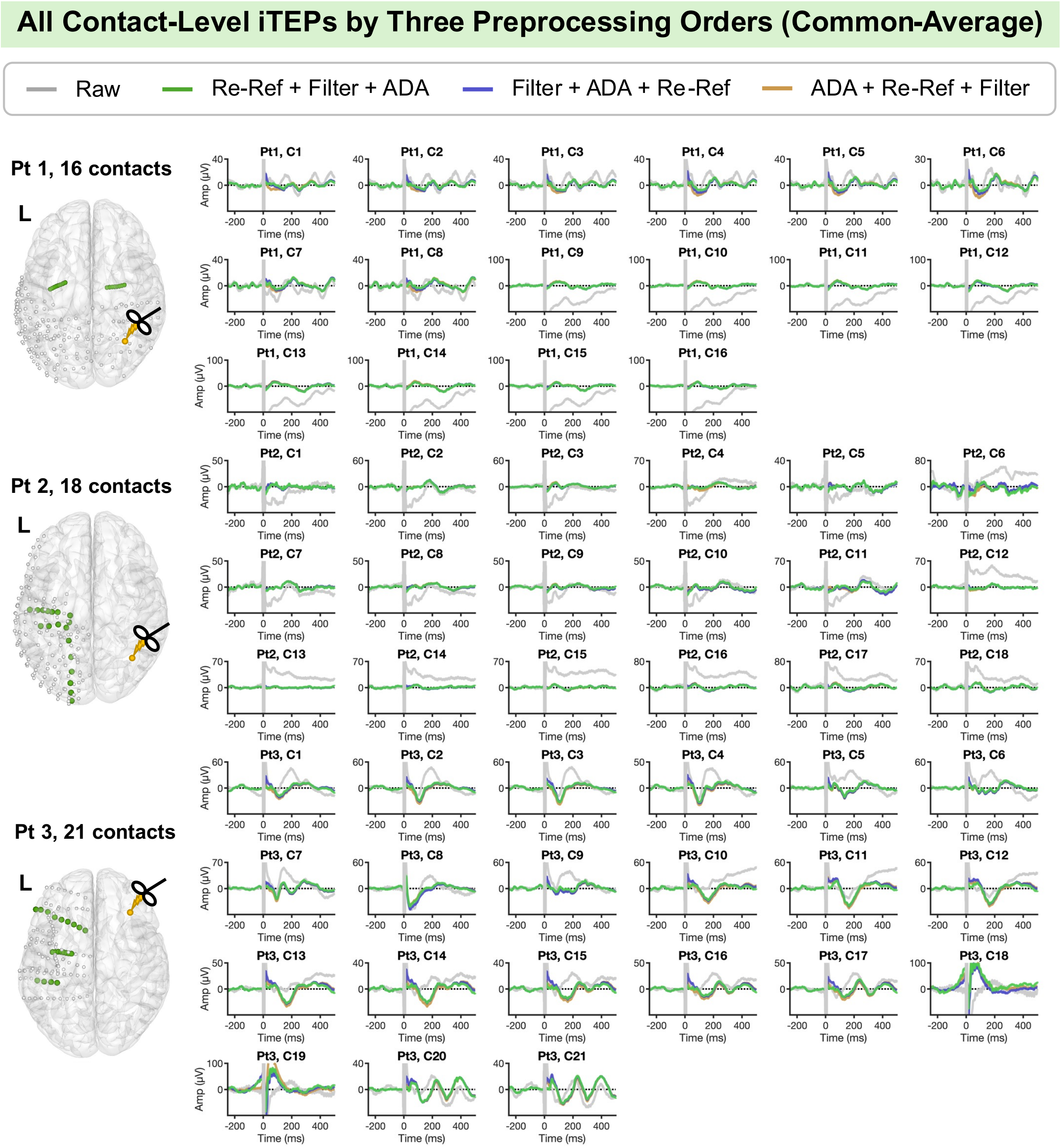

**Figure S13. Examples Illustrating the Effects of Artifact Window Selection on Trial-Level and Contact-Level iTEPs**

The resulting iTEPs using artifact windows of [–10, 15 ms] and [–10, 25 ms] were highly similar beyond the excluded window, with the signals within the 16–25 ms range preserved using the shorter window.

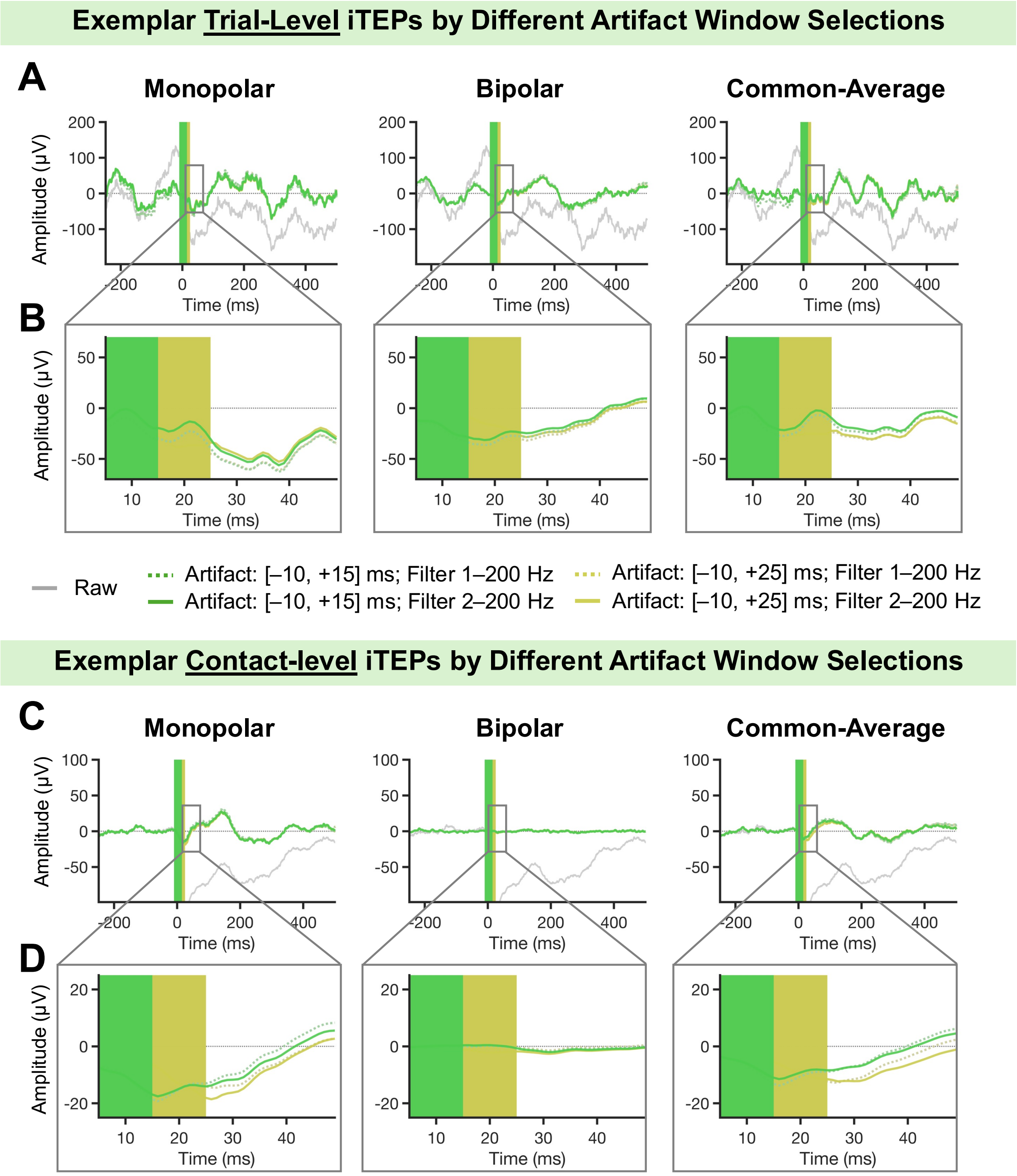

**Figure S14. Contact-level iTEPs by Different Selection of Artifact Windows (Monopolar Referencing)**

In most contacts, the resulting iTEPs using the two artifact windows were similar beyond the excluded window, with the signals in the 16–25 ms range preserved by the selection of a shorter window. Contacts showing noticeable differences (e.g., Pt 3-C18/19) were inherently noisy in the raw data (see Figure S10).

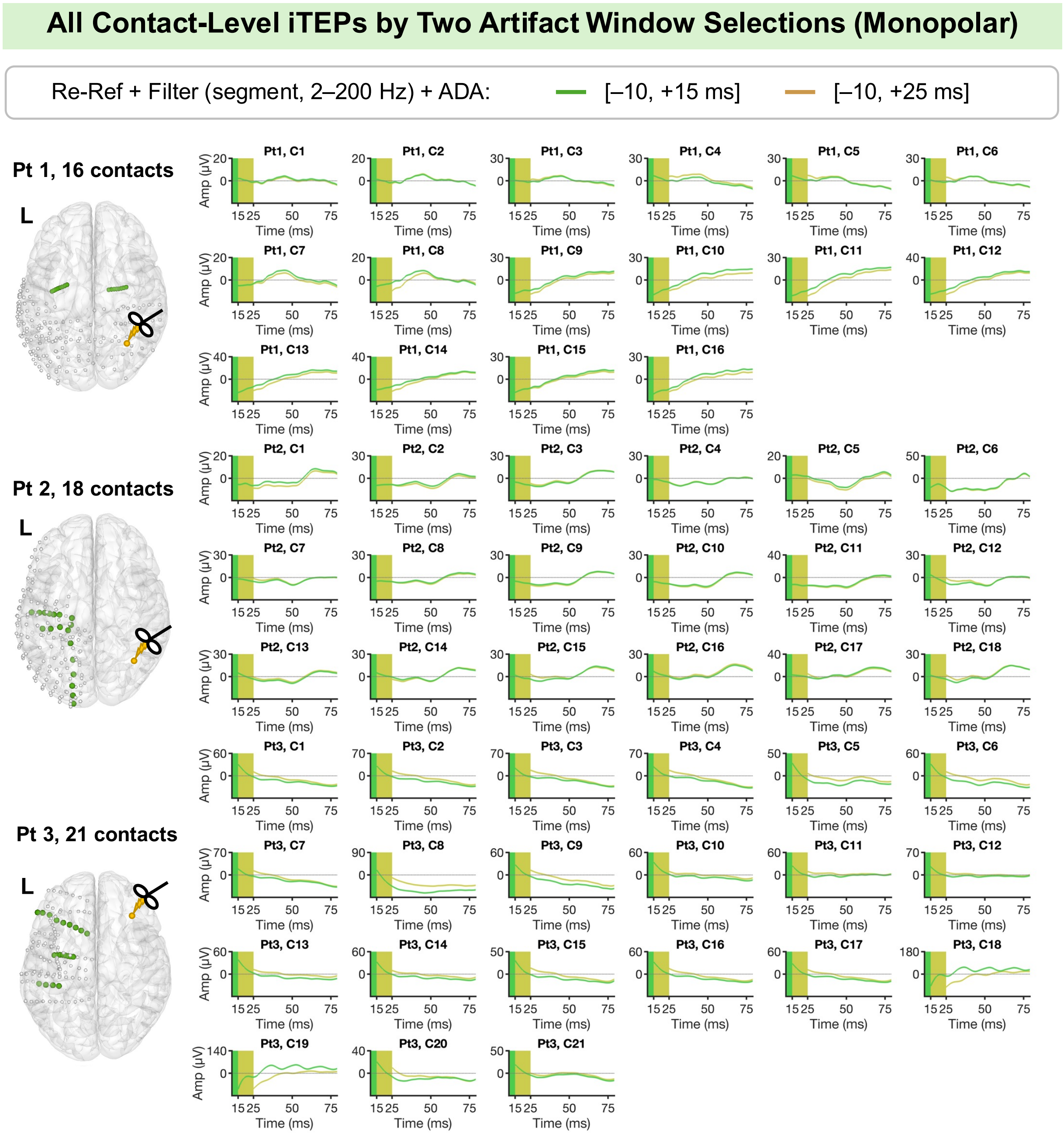

**Figure S15. Contact-level iTEPs by Different Selection of Artifact Windows (Bipolar Re-Referencing)**

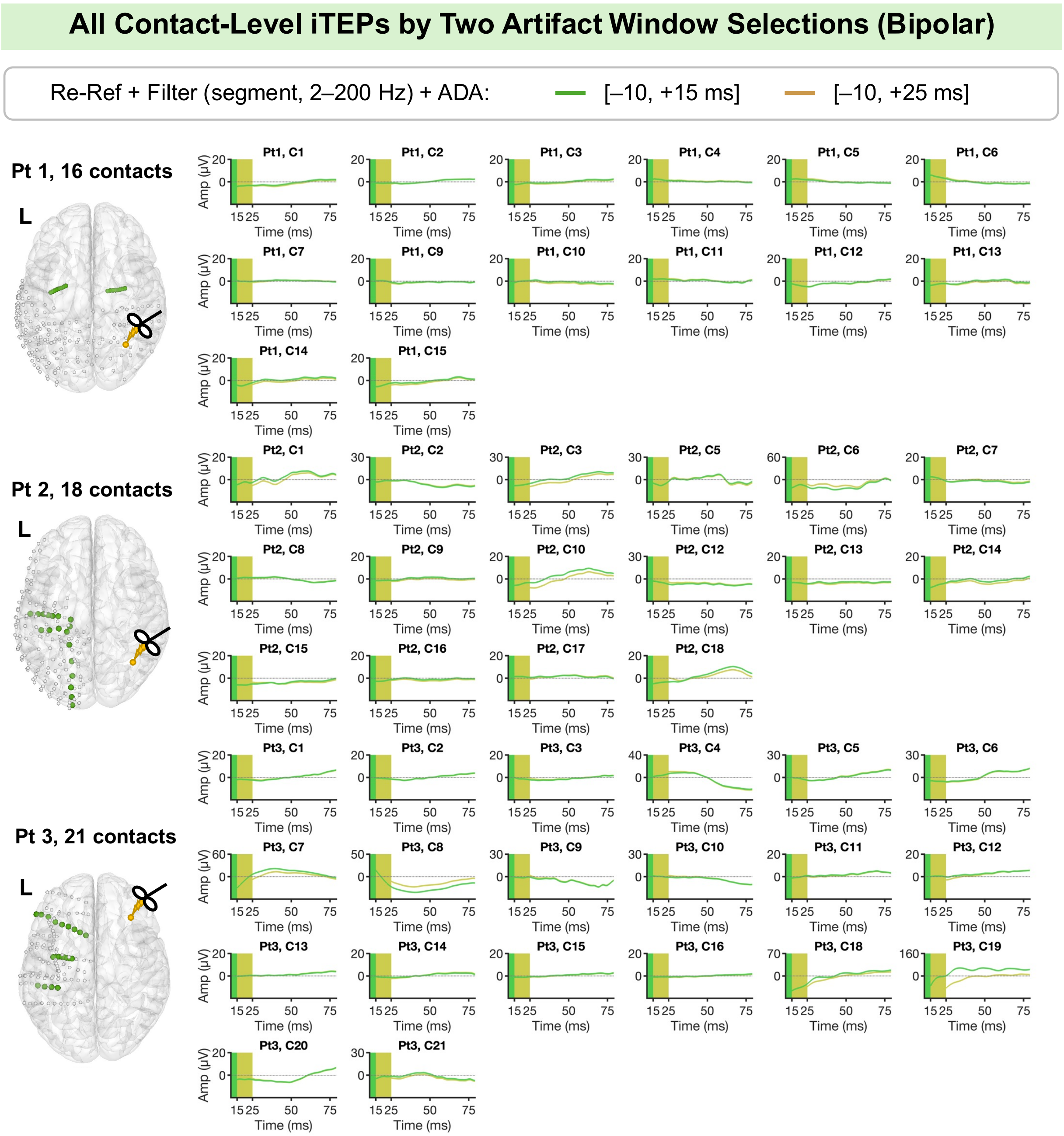

**Figure S16. Contact-level iTEPs by Different Selection of Artifact Windows (Common-Average Re-Referencing)**

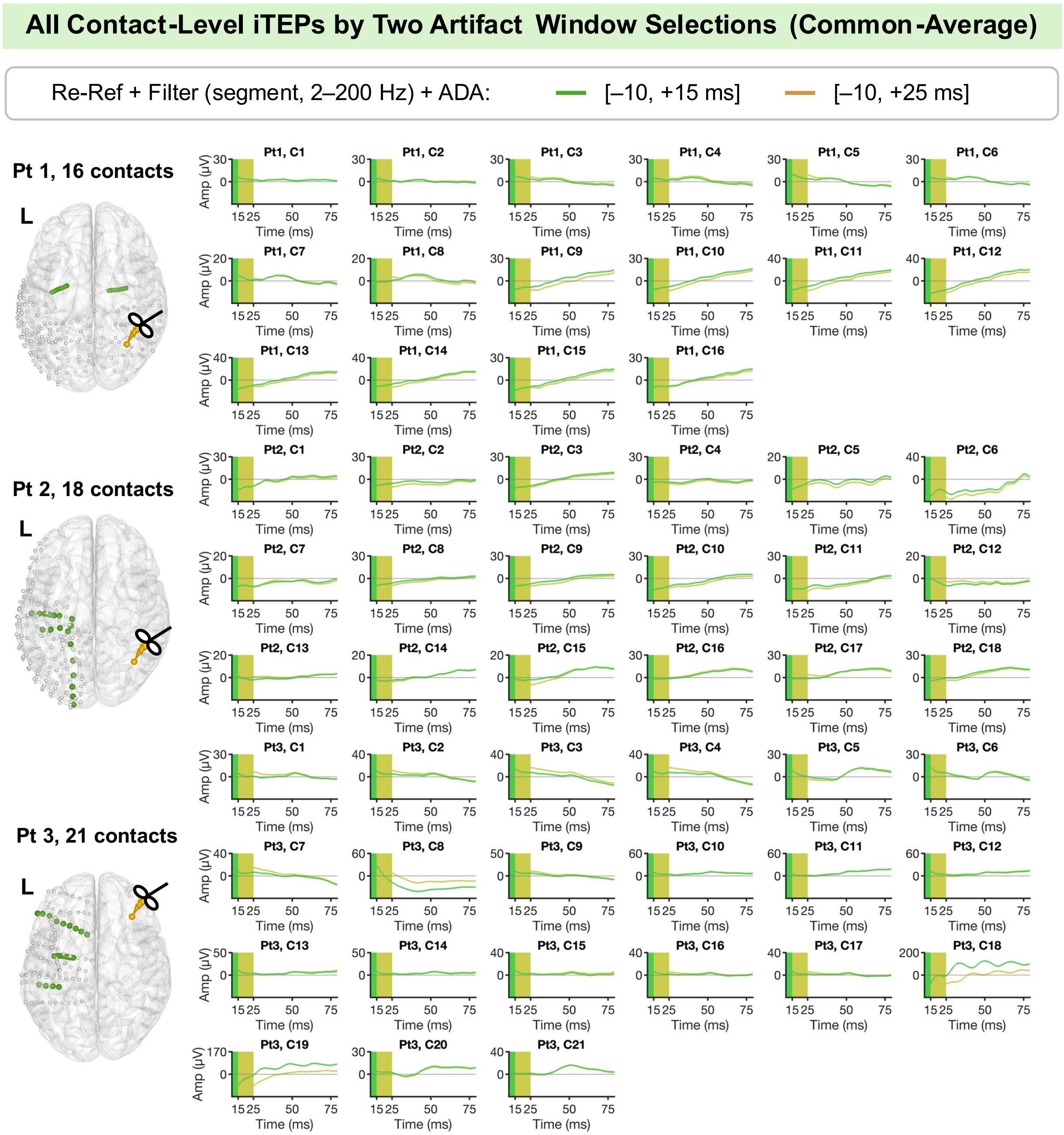

**Figure S17. Examples Illustrating the Utility of the Preprocessing Pipeline on ECoG-based Data**

On these two exemplar ECoG-based contacts, our preprocessing pipeline effectively mitigated artifacts and noise, yielding cleaner iTEPs at both trial and contact levels. Here, we applied monopolar and common-average referencing (excluding bipolar referencing, which is typically designed for sEEG) and tested two filtering strategies (segment-based vs. full-length-based filtering, both with 2–200 Hz bandpass filter). Consistent with results from sEEG data, segment-based filtering showed superior performance in avoiding filtering-induced distortions than full-length-based filtering, particularly in Contact 2.

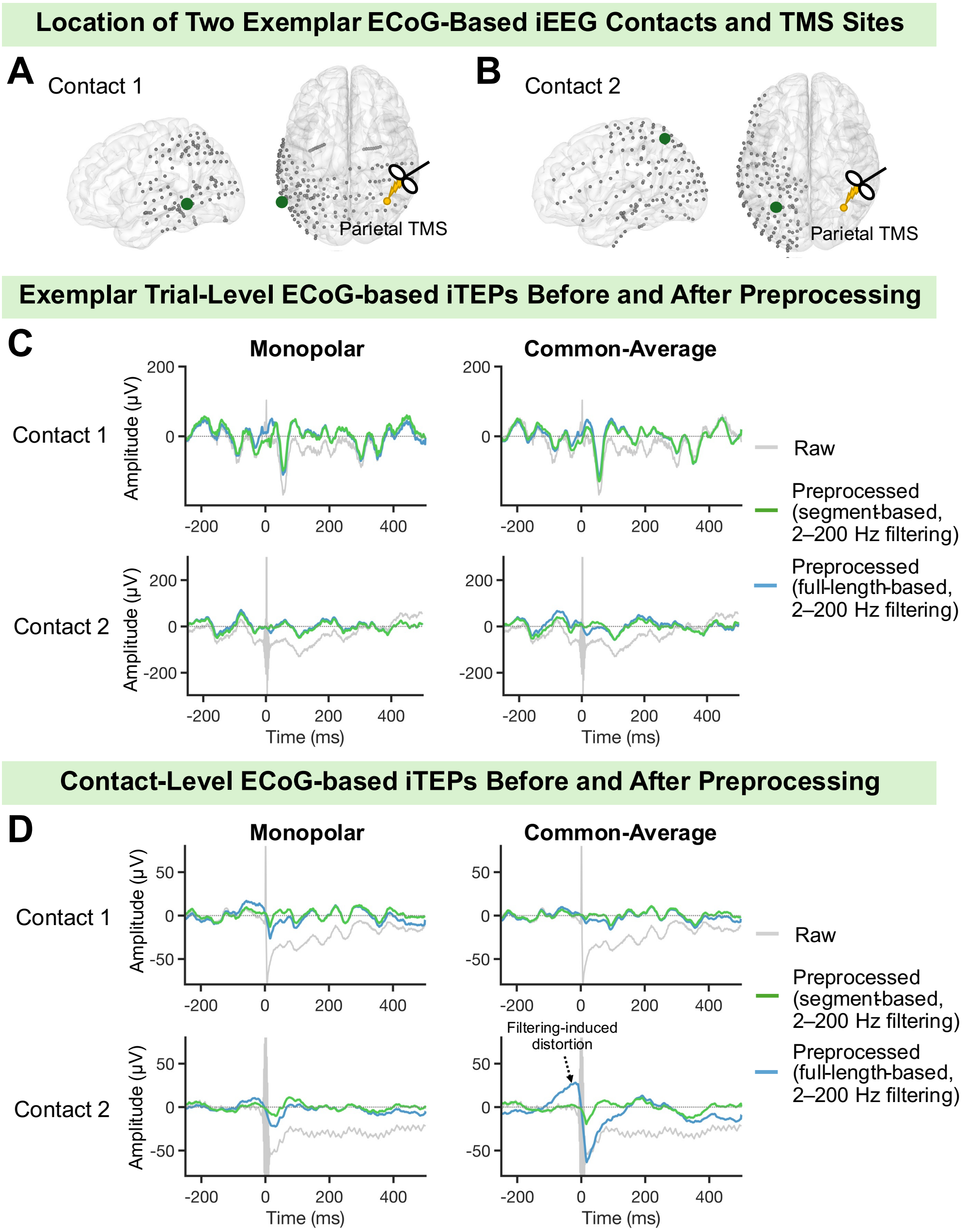
